# Intrinsic functional connectivity among memory networks does not predict individual differences in narrative recall

**DOI:** 10.1101/2023.08.31.555768

**Authors:** Kyle Kurkela, Maureen Ritchey

**Author notes:** Correspondence concerning this article should be addressed to: Maureen Ritchey, Boston College Department of Psychology and Neuroscience Chestnut Hill, MA.

## Abstract

Individuals differ greatly in their ability to remember the details of past events, yet little is known about the brain processes that explain such individual differences in a healthy young population. Previous research suggests that episodic memory relies on functional communication among ventral regions of the default mode network (“DMN-C”) that are strongly interconnected with the medial temporal lobes. In this study, we investigated whether the intrinsic functional connectivity of the DMN-C subnetwork is related to individual differences in memory ability, examining this relationship across 243 individuals (ages 18-50 years) from the openly available Cambridge Center for Aging and Neuroscience (Cam-CAN) dataset. We first estimated each participant’s whole-brain intrinsic functional brain connectivity by combining data from resting-state, movie-watching, and sensorimotor task scans to increase statistical power. We then examined whether intrinsic functional connectivity predicted performance on a narrative recall task. We found no evidence that functional connectivity of the DMN-C, with itself, with other related DMN subnetworks, or with the rest of the brain, was related to narrative recall. Exploratory connectome-based predictive modeling (CBPM) analyses of the entire connectome revealed a whole-brain multivariate pattern that predicted performance, although these changes were largely outside of known memory networks. These results add to emerging evidence suggesting that individual differences in memory cannot be easily explained by brain differences in areas typically associated with episodic memory function.

## 1 Introduction

Have you ever spoken with a family member about an event that happened long ago and admired the level of detail with which they could recall the event? While important differences exist at the extreme ends of the spectrum of memory ability, such as in patients with hippocampal damage (e.g., Patient H.M., see Corkin, 2002) or in individuals with superior autobiographical memory (LePort et al., 2012; Parker et al., 2006), experiences like these reinforce that there is substantial variation within the general population in the ability to recall the details of past events. Yet relatively little is known about the brain processes that explain such individual differences among healthy adults. In this study, we investigated the relationship between individual differences in memory performance and functional communication among regions known to be involved in episodic memory.

Prior studies examining the relationship between functional connectivity and individual differences in memory have largely focused on the functional connectivity of the hippocampus due to its important role in memory function (Bird & Burgess, 2008; Corkin, 2002; Squire, 1992). These studies have generally found that increased hippocampal functional connectivity is associated with better memory ability, including its bilateral functional connectivity (L. Wang, Negreira, et al., 2010) and its functional connectivity with cortical areas such as the lateral occipital cortex (Tambini et al., 2010) and the posterior medial cortex (Touroutoglou et al., 2015; L. Wang, LaViolette, et al., 2010). This positive relationship between hippocampal connectivity and memory ability can be interpreted as reflecting greater responsiveness of the hippocampus to time-varying signals across the brain in individuals with better memory, or conversely, disruption in hippocampal communication in individuals with worse memory. However, these studies have often been underpowered, both in terms of the number of participants collected (Marek et al., 2022) and in terms of the amount of data collected per participant (Anderson et al., 2011; Gordon, Laumann, Gilmore, et al., 2017; Laumann et al., 2015). Brain-wide association studies, for example, require thousands of participants to achieve acceptable levels of statistical power (Marek et al., 2022), though this problem is mitigated by using a region of interest-based approach to reduce the number of comparisons or by focusing on multivariate patterns within the connectome. Functional connectivity studies may additionally benefit from more data being collected per participant, as it has been recently shown that at least 30 minutes of high quality MRI data is required to achieve good levels of reliability of the functional connectome (i.e., r > 0.85; Gordon, Laumann, Gilmore, et al., 2017). This is important because the reliability of a measurement places a key constraint on the measurable effect size of the correlation between two constructs of interest (e.g., functional connectivity and behavior). Most existing studies relating functional connectivity to memory ability have typically used data from one MRI scan, often comprising 6-8 minutes of data.

Because prior research has largely focused on the functional connectivity of the hippocampus, relatively less is known about how functional connectivity among cortical networks relates to episodic memory. Episodic memory has been linked to the functions of the default mode network (DMN), a set of brain regions that tend to be co-activated during rest and during tasks involving episodic construction (Buckner et al., 2008; Buckner & DiNicola, 2019; Ritchey & Cooper, 2020). In particular, task based MRI studies have shown that memory tasks recruit a ventral subnetwork of the DMN that is strongly interconnected with the medial temporal lobes (Andrews-Hanna et al., 2010; Barnett et al., 2021; Buckner & DiNicola, 2019). This DMN subnetwork, labeled DMN-C in recent brain parcellations (Schaefer et al., 2018; Yeo et al., 2011), consists of the retrosplenial cortex, parahippocampal cortex, and the posterior angular gyrus. The DMN-C subnetwork is commonly co-activated with an adjacent, more dorsal DMN subnetwork, labeled DMN-A, which consists of medial frontal and parietal regions. Recent work has shown that these two subnetworks are dissociable in terms of their functional connectivity during event perception (Cooper et al., 2021) as well as their contributions to memory retrieval (Kurkela et al., 2022). In the latter study, when activity estimates from both ventral and dorsal DMN regions were included in a model predicting retrieval success, only the ventral regions significantly predicted success in retrieving event-specific associations. Based on these findings, we hypothesize that the function of the DMN-C subnetwork may be central in determining individual differences in episodic memory ability, and in particular, the ability to recall specific details associated with past events.

More recent experiments have looked at the relationship between memory and intrinsic functional connectivity using larger samples (Przeździk et al., 2019) or incorporating a broader set of brain regions (Sneve et al., 2017; van Buuren et al., 2019). These studies suggest that there is a complex, distributed pattern of whole brain functional connectivity beyond the hippocampus that is related to individual differences in episodic memory ability, but the exact nature of this relationship has varied from study to study. Some studies suggests that superior rememberers have default mode networks that are decoupled from perceptual regions of the brain (Sneve et al., 2017), whereas others suggest that superior rememberers are characterized by decreased connectivity within the ventral default mode network and strong connectivity between the dorsal default mode network and the frontal-parietal control network (van Buuren et al., 2019). Other work suggests that memory ability is associated with hippocampal-cortical connectivity patterns that gradually change along the hippocampal long axis (Przeździk et al., 2019). Taken together, these studies suggest that there is likely some whole brain functional connectivity pattern that is related to episodic memory ability, but the exact nature of this whole brain pattern remains unclear.

In the present study, we investigated the relationship between intrinsic functional connectivity and episodic memory ability, focusing on narrative recall as a hallmark of intact episodic memory that has been linked specifically to the function of the DMN (Lee et al., 2020; Ritchey & Cooper, 2020). This study improved on previous work by incorporating a relatively large sample size and by incorporating measures of “generalized functional connectivity” that increase the amount of functional data included per participant, which improves the reliability of functional connectivity estimates and may enhance prediction of cognitive abilities (Elliott et al., 2019). We took two complementary approaches to examining this brain-behavior relationship. First, we took a hypothesis-driven approach to test whether functional connectivity of the DMN-C subnetwork was specifically related to individual differences in narrative recall. We complemented these analyses with a data-driven approach using connectome-based predictive modeling (Shen et al., 2017) to determine if there were any patterns of whole-brain connectivity that were predictive of narrative recall ability.

## 2 Methods

To answer our research questions, we analyzed data from the Cambridge Center for Aging and Neuroscience (Cam-CAN) repository (Shafto et al. 2014; Taylor et al. 2017). The Cam-CAN repository is a large-scale, cross-sectional, openly available cognitive neuroscience dataset collected by the University of Cambridge. Below we summarize the key characteristics of the Cam-CAN dataset, focusing specifically on the subsets of the dataset that were utilized in the present report. Our analysis plan was preregistered on the Open Science Framework: https://osf.io/9xcu3/?view_only=1ac6856b773249cfa0767dd3d005a9ae.

### 2.1 Participants

Participants included 243 participants between the ages of 18 and 50 sampled from the original set of 653 Cam-CAN participants who had data available at the time of our access. Participants in the original set were equally sampled from each decile of age from 18-87 years of age with approximately equal numbers of men and women in each decile. They were required to be cognitively healthy, to not have a serious psychiatric condition, to have met hearing and English language requirements for experiment participation, and to be eligible for MRI scanning (Shafto et al., 2014). Participants gave informed consent prior to their participation in the study, as described in Shafto et al. (2014). Of the 653 available subjects, 7 were missing at least one of the five MRI scans (*see MRI Data*), 1 was missing data from logical memory subtest of the Wechsler Memory Scale, 19 were missing Cattell Fluid Intelligence scores, and 1 subject was missing ACE-R data (*see Behavioral Data*). To deal with missing data, we took a listwise deletion approach such that if a participant was missing any of the variables of interest, they were removed from further analysis. Participants were excluded from the current set of analyses if they met the following exclusion criteria. First, participants were excluded from the analysis if they had 2 or more functional runs that had a mean framewise displacement greater than 0.3mm, to mitigate the effects of motion on functional connectivity estimates (n = 217). Second, participants were excluded from the analysis if they were older than 50 years of age to mitigate the influence of advanced aging on our results (n = 185). After applying these additional exclusion criteria, we were left with 243 subjects to analyze. These 243 subjects were on average 36.28 years old (median = 36.33 years, SD = 8.42) and 122 self-reported as female and 121 as male.

### 2.2 MRI Data

The subset of the Cam-CAN dataset analyzed in the present report contained a single high resolution T1-weighted anatomical scan, three functional scans, and a single field map to correct for magnetic field inhomogeneities. The three functional scans included a movie-watching scan, a resting-state scan, and a sensorimotor scan (detailed below). The anatomical and field map images were used during the preprocessing of the functional scans. The three functional scans were used to estimate each subject’s intrinsic functional connectome (see *Functional Connectivity*). The anatomical scan was acquired using a Magnetization Prepared RApid Gradient Echo (MPRAGE) sequence with the following parameters: Repetition Time (TR) = 2250 ms; Echo Time (TE) = 2.99 ms; Inversion Time (TI) = 900 ms; flip angle = 9 degrees; field of view (FOV) = 256mm x 240mm x 192mm; voxel size =1mm isotropic; GRAPPA acceleration factor = 2; acquisition time of 4 min and 32 sec. The movie watching scan involved participants watching an edited version of Alfred Hitchcock’s movie “Bang, You’re Dead”. A total of 193 volumes were acquired using a multi-echo, T2*-weighted EPI sequence (TR = 2470 milliseconds, five echoes [TE = 9.4 milliseconds, 21.2 milliseconds, 33 milliseconds, 45 milliseconds, 57 milliseconds], flip angle = 78 degrees, 32 axial slices of thickness of 3.7 mm with an interslice gap of 20%, FOV = 192mm × 192 mm, voxel-size = 3 mm × 3mm × 4.44 mm) with an acquisition time of 8 minutes and 13 seconds. The resting-state scan involved participants resting in the scanner with their eyes closed. During the sensorimotor scan, participants were presented with visual checkerboards and auditory tones, either in isolation or simultaneously. They were instructed to respond with a button press when they were presented with any stimuli (either visual, auditory, or both visual and auditory). The resting state and sensorimotor scans had the same scanning parameters: a total of 261 volumes were acquired, each containing 32 axial slices acquired in descending order, slice thickness of 3.7 mm with an interslice gap of 20%; TR = 1970 ms; TE = 30 ms; flip angle = 78 degrees; FOV = 192 mm × 192 mm; voxel-size = 3 mm × 3 mm × 4.44 mm) and an acquisition time of 8 min and 40 sec. The fieldmap consisted of an SPGR gradient-echo sequence with the same parameters as the resting state and sensorimotor tasks, but with two TEs (5.19 ms and 7.65 ms).

### 2.3 Regions of Interest

Regions of interest (ROIs) were taken from the Schaefer cortical parcellation (Schaefer et al., 2018). Specifically, we used the 400-area resolution, 17-network parcellation. We focused our analyses *a-priori* on three sets of parcels from this atlas: DMN-C regions (number of parcels: left hemisphere = 7, right hemisphere = 6), DMN-A regions (number of parcels: left hemisphere = 18, right hemisphere = 16), and all other regions (number of parcels: left hemisphere = 176, right hemisphere = 178). DMN-C regions consisted of the bilateral parahippocampal gyrus, bilateral retrosplenial cortex, and the bilateral posterior angular gyrus (see Figure 1a, blue regions). DMN-A regions included the bilateral posterior cingulate cortex, the bilateral precuneus, the bilateral medial prefrontal cortex, the bilateral anterior angular gyrus, and a bilateral section of the dorsal prefrontal cortex (see Figure 1a, yellow regions). We focused on regions in the DMN-C and DMN-A networks due to their roles in episodic memory and simulation (Buckner & DiNicola, 2019; Ritchey & Cooper, 2020) and their correspondence with ventral and dorsal subnetworks we have previously identified (Cooper et al., 2021; Kurkela et al., 2022). We note that these were inadvertently mislabeled in our pre-registration. To supplement this cortical atlas, we included hippocampal ROIs created using the anatomical delineations from Ritchey and colleagues (2015). Specifically, we used hippocampal ROIs comprising the hippocampal head, the hippocampal body, and the hippocampal tail from the right and left hemispheres. These six hippocampal ROIs were added to the 400 cortical ROIs to form the functional connectome (i.e., 406x406 ROI-to-ROI connectivity matrix).

**Figure 1.**
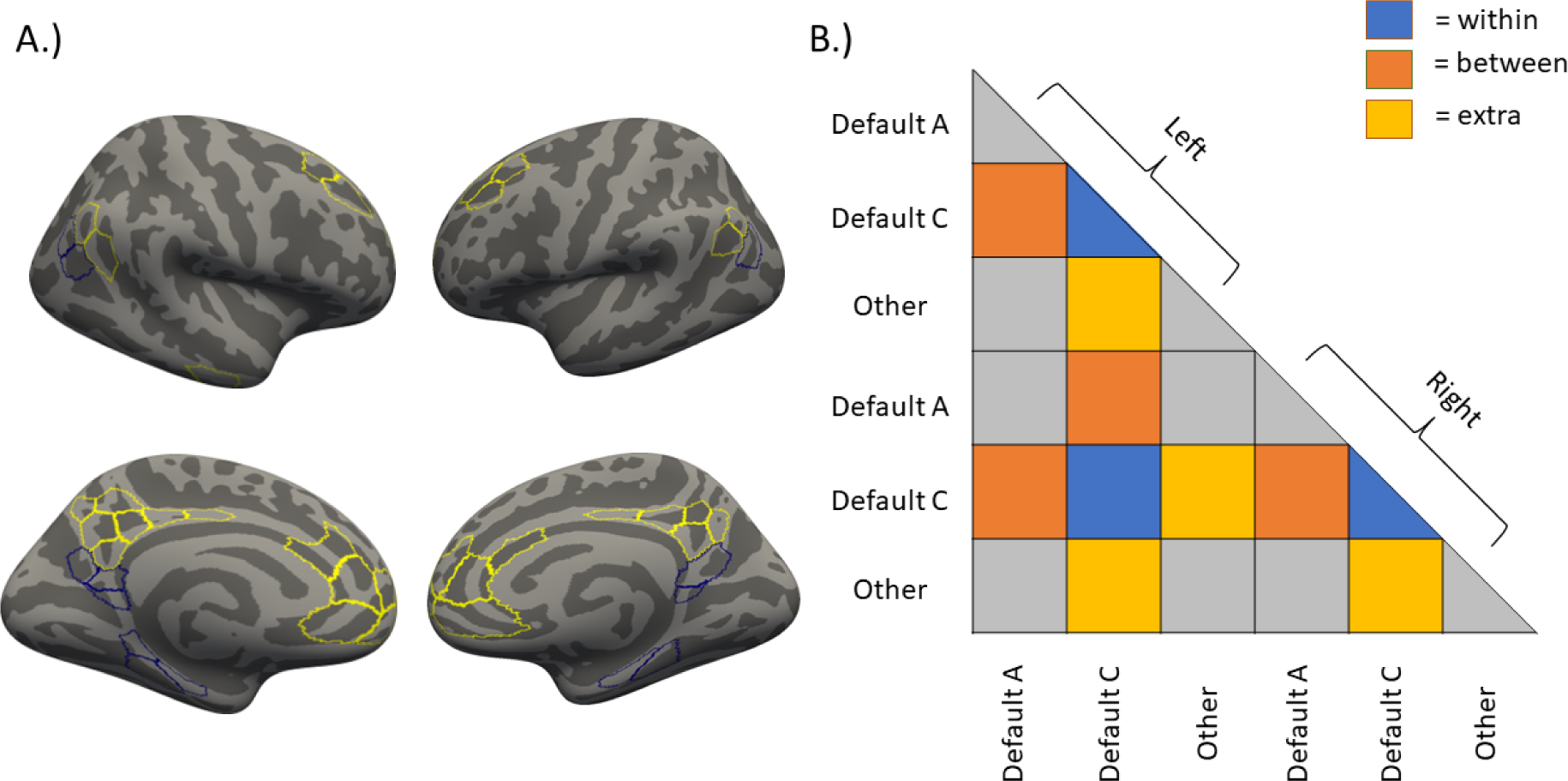
Regions and connections of interest. A.) Regions of interest were based on the 400 parcel 17-network parcellation from the Schaefter atlas (Schaefer et al., 2018). Here we highlight the DMN-C (blue) and DMN-A (yellow) networks from this atlas. This visualization was created in freeview using the *fsaverage* inflated surface template provided in the Schaefer et al. (2018) atlas. B.) The lower triangle of the functional connectome. Connections of interest are highlighted, including “within” connections (blue), “between” connections (orange), and “extra” network connections (yellow) of the DMN-C. Connections of no interest are shown in gray.

### 2.4 Behavioral Data

Summary statistics for the behavioral and neural variables of interest are presented in Table 1. Behavioral data included performance on the following cognitive assessments: the logical memory subtest from the Wechsler Memory Scale Third UK edition (Weschler, 1999), the Cam-CAN emotional memory task (Taylor et al., 2017), the Addenbrook Cognitive Examination-Revised (variable name: ACER), and the Cattell Test of Fluid Intelligence (variable name: Cattell). The logical memory subtest from Wechsler Memory Scale involved having participants read two short passages and subsequently verbally recall as many story details as possible at two different time points: first immediately after reading the short passages and then again after a ∼20 min delay. Verbal recalls were scored for the number of story details correctly recalled at each point in time. The number of story details recalled at both points in time were averaged together to form a narrative recall score, which serves as our primary measure of memory ability and primary dependent variable. The emotional memory task included recall of object-scene associations (see Taylor et al., 2017 for a description of this task), and the total number of detailed scene recalls was incorporated here as an additional measure of memory ability. Narrative recall and emotional memory scores were standardized across the sample and averaged to create a composite measure of memory ability. Analyses additionally incorporated two general measures of cognitive function. The ACER is a standardized cognitive battery originally designed for dementia screening. The battery is designed to test participants’ ability in 5 different cognitive domains, including attention, memory, fluency, language, and visual-spatial. The total score on this battery was used as an index of general cognitive function. The Cattell Test of Fluid Intelligence is a timed pen and paper task where participants are required to solve a series of non-verbal puzzles. Here we use the total score on this task as a general index of fluid intelligence.

**Table 1:** Data Summary. Statistical summary of data analyzed for the current report. *mem-narr* = number of story details recalled in the logical memory subtest of the Wechsler Memory Scale – average of immediate and delayed, *mem-emo* = total number of detailed scene recalls in the Cam-CAN emotional memory task, *Cattell* = total score on the Cattell Test of Fluid Intelligence, *ACER* = total score on Addenbrook’s Cognitive Evaluation Revised (ACE-R), *fd* = mean framewise displacement, *within* = average functional connectivity estimate (Pearson’s correlation *r*) of DMN-C—DMN-C connections, *between* = average functional connectivity estimate (Pearson’s correlation *r*) of DMN-C–DMN-A connections, *extra* = average functional connectivity estimate (Pearson’s correlation *r*) for DMN-C–rest of the brain connections, n = number of complete observations. Correlations were calculated using all available complete pairs of data. All values are rounded to two decimal places where appropriate. ^a^ Sex was coded such that Female = 1, Male = 0. ^b^ Half of subjects completed Cam-CAN’s emotional memory task (see Shafto et al. 2014). ^c^ 8 Subjects Missing Cattell Scores.

| Variable | mean | min | max | sd | n | Correlations |  |  |  |  |  |  |  |  |  |
| --- | --- | --- | --- | --- | --- | --- | --- | --- | --- | --- | --- | --- | --- | --- | --- |
|  |  |  |  |  |  | mem-narr | mem-emo | Age | Sex | Cattell | ACER | fd | within | between | extra |
| mem-narr | 15.28 | 2.50 | 23 | 3.50 | 243 |  |  |  |  |  |  |  |  |  |  |
| mem-emo | 52.98 | 4 | 105 | 23.98 | 120<sup>b</sup> | 0.44 |  |  |  |  |  |  |  |  |  |
| Age | 36.28 | 18.50 | 49.83 | 8.42 | 243 | -0.13 | -0.32 |  |  |  |  |  |  |  |  |
| Sex<sup>a</sup> | 0.50 | 0 | 1 |  | 243 | 0.14 | 0.10 | -0.07 |  |  |  |  |  |  |  |
| Cattell | 36.35 | 22 | 44 | 4.23 | 235<sup>c</sup> | 0.35 | 0.30 | -0.25 | -0.19 |  |  |  |  |  |  |
| ACER | 96.54 | 74 | 100 | 3.63 | 243 | 0.40 | 0.31 | 0.12 | 0.03 | 0.39 |  |  |  |  |  |
| fd | 0.14 | 0.05 | 0.25 | 0.04 | 243 | -0.10 | -0.11 | 0.23 | 0.02 | -0.22 | -0.14 |  |  |  |  |
| within | 0.31 | 0.17 | 0.50 | 0.06 | 243 | 0.02 | 0.09 | -0.21 | 0.02 | 0.09 | 0.04 | -0.23 |  |  |  |
| between | 0.15 | 0.03 | 0.26 | 0.05 | 243 | 0.04 | -0.08 | -0.01 | 0.09 | -0.06 | 0.03 | -0.10 | 0.53 |  |  |
| extra | -0.02 | -0.06 | 0.02 | 0.01 | 243 | -0.09 | -0.16 | 0.14 | 0.02 | -0.06 | -0.07 | 0.14 | -0.55 | -0.63 |  |
| hipp | 0.00 | -0.03 | 0.04 | 0.01 | 243 | -0.10 | -0.22 | -0.02 | -0.18 | 0.01 | -0.03 | -0.07 | 0.02 | -0.19 | 0.28 |

### 2.5 Analysis

#### 2.5.1 MRI Quality Control

The quality of the functional MRI data was assessed using the *MRIQC* software package (Esteban et al., 2017). Quality reports generated from this software package were visually inspected for scanner artifacts and motion-related corruption. All scans that had a mean framewise displacement greater than 0.3mm were excluded from further analysis. If subjects had 2 or more scans with a mean framewise displacement greater than 0.3mm, then the subject was excluded from further analysis (see *Participants*). This conservative approach was adopted in order to limit the effects of head motion on functional connectivity measures (see Cooper et al., 2021 for a similar exclusion criterion; Power et al., 2012, 2014).

#### 2.5.2 MRI Preprocessing

MRI data were preprocessed using *fMRIPrep* 20.2.0 (Esteban et al., 2018, 2019). Processing steps for the T1w images included correction for intensity non-uniformity, skull stripping, brain tissue segmentation, and volume-based spatial normalization to the ICBM 152 Nonlinear Asymmetrical template version 2009c. Processing steps for the 3 BOLD runs included slice-timing correction, realignment, using the fieldmap to optimize co-registration of the functional images to the anatomical reference image, normalization of the BOLD images to the ICBM 152 Nonlinear Asymmetrical template version 2009c, and the calculation of confounding time-series including basic 6 head-motion parameters (x,y,z translation; pitch, roll, yaw rotation), temporal derivatives and the quadratic terms of the head-motion parameters, noise components from a principal components analysis based denoising routine (*CompCorr*), framewise displacement, and DVARS. For a much more detailed description of the processing pipeline please refer to the Supplemental Materials which contains the recommended *fMRIPrep* boilerplate.

#### 2.5.3 Functional Connectivity

Functional connectivity analyses were performed using the *CONN* toolbox (Whitfield-Gabrieli & Nieto-Castanon, 2012). Confounds removed from each voxel’s time series included six head motion parameters and their temporal derivatives, up to the first six *aCompCor* components from a combined WM and CSF mask, framewise displacement, and the global signal^1^ as calculated by *fMRIPrep*. Additional spike regressors were included for any time points that exceeded a FD of 0.6 mm and/or a standardized DVARS of 2. After regression of motion confounds, BOLD data was band-pass filtered with a high-pass filter of 0.008 Hz and a low-pass filter of 0.1 Hz. No additional modeling was performed for the movie-watching and the resting state data. For the sensorimotor task, task-related activations were regressed out of the time series prior to calculation of functional connectivity. Specifically, task related activation was modeled in the sensorimotor task by convolving stick functions placed at stimulus onsets with SPM12’s hemodynamic response function. ROI-to-ROI functional connectivity was calculated using the Pearson’s correlation coefficient after denoising and task modeling. The ROI-to-ROI intrinsic functional connectivity estimates were created by averaging functional connectomes calculated from all available functional scans, resulting in a measure of “generalized” functional connectivity intrinsic to each individual (Elliott et al., 2019). The resulting ROI-to-ROI intrinsic functional connectomes were then summarized to three terms to test our research hypotheses. *Within* subnetwork connectivity was operationalized as the average of all functional connections among DMN-C nodes in the Schaefer et al. (2018) atlas, *between* subnetwork connectivity as the average connectivity between all DMN-C and DMN-A nodes, and *extra* network connectivity as the average of all connections between the DMN-C nodes and all other nodes not contained in DMN-C or DMN-A. See Figure 1b for an illustration. Although averaging across the three scan types has the advantage of increasing the amount of fMRI data used to estimate functional connectivity and increasing the reliability of the resulting functional connectivity estimates (Elliott et al., 2019), it may obscure task-specific relationships between functional connectivity and behavior, particularly in the context of tasks that vary considerably in their content and demands. Thus, we additionally conducted exploratory analyses in which functional connectivity relationships were analyzed separately for each of the three scan types (resting-state, movie-watching, and sensorimotor task).

#### 2.5.4 Statistical Modeling

All statistical models, tables, and figures were created using *R (R Core Team, 2022)*. Linear regression models were used to estimate thes relationship between summary measures of functional connectivity and memory ability, on their own and then including covariates for age, sex, average framewise displacement, fluid intelligence, and cognitive function as defined under *Behavioral Data*. Standardized beta coefficients are reported. Summary tables for the linear regressions were created using a combination of the R packages *gtsummary (Sjoberg et al., 2021)* and *flextable (Gohel & Skintzos, 2023)*. All figures were created using the R package *ggplot2 (Wickham, 2016)*.

#### 2.5.5 Connectome Based Predictive Modeling

To complement the hypothesis driven approach described above, the current report used connectomic based predictive modeling (CBPM; Shen et al., 2017). CBPM involves calculating a functional connectome for each individual and determining which connections are statistically related to a behavioral variable of interest. All connections that are related to behavior above some arbitrarily defined threshold (e.g., at *p* < .01) are then summarized by separating out connections with a significant positive correlation with behavior from those that have a significant negative correlation with behavior. The connections in the connectome with significant positive and negative correlations with behavior are then summed into separate positive and negative terms for use in a linear regression predicting behavior. The analysis method then estimates a linear regression using these summed positive and negative connection terms to predict the behavior of a left-out participant in a leave-one-out cross validation procedure. Successful prediction of a behavioral variable using the functional connectome is then determined by correlating the predicted behavioral scores for each participant with the observed behavioral scores with statistical significance determined using a null permutation procedure that reruns the entire analysis a given number of times, each time randomly pairing connectomes with behavioral scores. All CBPM analyses were performed using a connection selection threshold of *p* < .01 and the null distribution was formed from 100 null simulations. The results from alternative connection selection thresholds are reported in the Supplemental Materials. Statistical significance of the CBPM analysis was determined by estimating the proportion of null simulations that resulted in better predictive performance than the actual analysis. All CBPM analyses were performed using modified analysis code published by Shen and colleagues (2017). To control for nuisance variables, the CBPM analysis code published by Shen and colleagues was modified to use the MATLAB function *partialcorr* when selecting connections used for prediction. To determine which connections in the connectome the models were relying on to make their predictions, we also performed computational lesion analyses. In a computational lesion analysis, a predictive model is iteratively fit while excluding a set of features. If model performance is hindered by the removal of a set of features, then it is inferred that the model is relying on these features to make successful out-of-sample predictions.

## 3 Results

### Functional connectivity within the Default-C subnetwork

Average intrinsic functional connectivity among DMN-C regions was not related to narrative recall (*β* = 0.06, SE = 0.23, *t*(241) = 0.28, *p* = 0.78; see Figure 2a). This pattern did not change after controlling for age, sex, and average framewise displacement (*β* = −0.10, *SE* = 0.23, *t*(238) = −0.42, *p* = 0.68) or after also controlling for fluid intelligence and cognitive function (*β* = −0.06, *SE* = 0.21, *t*(228) = −0.28, *p* = 0.78).

**Figure 2:**
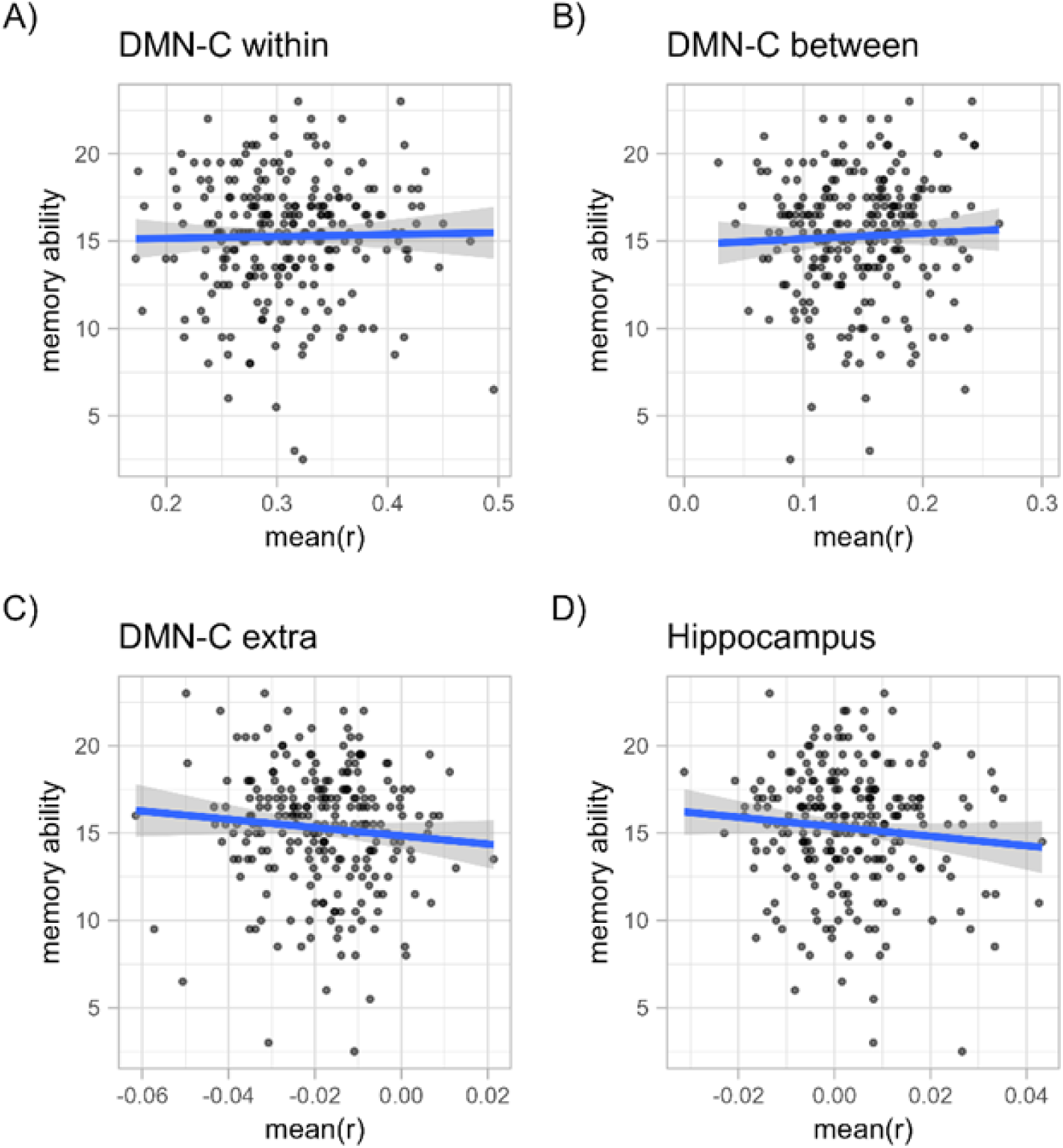
Relationships between memory ability and intrinsic functional connectivity among memory networks. Scatter plots and best fit linear regression line of narrative recall performance on average intrinsic connection strength A) among DMN-C regions, B) between DMN-C and DMN-A regions, C) DMN-C and all other regions, and D) the Hippocampus and all regions. There was little evidence to suggest that the strength of these connections was predictive of individual differences in memory ability.

### Functional connectivity between the Default C and Default A subnetworks

Average between subnetwork connectivity did not predict narrative recall on its own (*β* = 0.15, SE = 0.23, *t*(241) = 0.66, *p* = 0.51; see Figure 2b), nor did it predict narrative recall when controlling for age, sex, and average framewise displacement (*β* = 0.07, SE = 0.22, *t*(238) = 0.33, *p* = 0.74), nor when additionally controlling for fluid intelligence and cognitive function (*β* = 0.19, SE = 0.20, *t*(228) = 0.96, *p* = 0.34).

### Functional connectivity between the Default C subnetwork and all other brain regions

The average strength of functional connections from DMN-C regions to regions in networks other than the DMN-A and DMN-C did not predict narrative recall on its own (*β* = −0.31, *SE* = 0.22, *t*(241) = −1.39, *p* = 0.17; see Figure 2c), when controlling for age, sex, and average framewise displacement (*β* = −0.24, *SE* = 0.23, *t*(238) = −1.05, *p* = 0.29), nor when additionally controlling for fluid intelligence and cognitive function (*β* = −0.25, *SE* = 0.20, *t*(228) = −1.23, *p* = 0.22).

### Functional connectivity of the hippocampus

Although not part of our preregistered set of analyses, we decided to look at the relationship between narrative recall and intrinsic functional connectivity of the hippocampus given prior research linking this region to individual differences in memory (Touroutoglou et al., 2015; L. Wang, LaViolette, et al., 2010; L. Wang, Negreira, et al., 2010). We set about testing this in a similar manner to our preregistered set of analyses. We found that the average strength of all hippocampal connections was not a significant predictor of narrative recall on its own (*β* = −0.34, *SE* = 0.22, *t*(241) = −1.5, *p* = 0.13; see Figure 2d). This result held when we statistically controlled for age, sex, and average framewise displacement (*β* = −0.29, *SE* = 0.23, *t*(238) = −1.28, *p* = 0.2), and when we further controlled for fluid intelligence and cognitive function (*β* = −0.26, *SE* = 0.20, *t*(228) = −1.3, *p* = 0.2). We note here that the inclusion of global signal regression in our analysis pipeline influenced this result in that exclusion of this step resulted in a significant negative association between hippocampal connectivity and narrative recall (see Supplemental Materials for further discussion).

### Task-specific relationships between functional connectivity and memory ability

To explore whether there might be task-specific relationships between functional connectivity and memory, we re-ran the above analyses separately for each scan type (resting-state, movie-watching, and sensorimotor task). There were no significant relationships between any of the task-specific functional connectivity measures and narrative recall, all *p*s > .05.

We additionally sought to determine whether this pattern of null results was specific to the relationship between functional connectivity and narrative recall performance. To do so, we calculated a composite memory ability score incorporating the narrative recall measure as well as performance on an emotional memory task collected in a subset of our sample (N=120). This composite measure was included as the dependent variable in the regression models analogous to those reported above. The results were largely similar to those reported for the narrative recall analyses above, with no significant relationships between composite memory ability and functional connectivity among DMN regions. One exception was that in these analyses, there was a significant negative relationship between hippocampal functional connectivity and composite memory ability, *β* = −0.21, *SE* = 0.08, *t*(118) = −2.73, *p* = 0.007, which held up after controlling for age, sex, framewise displacement, fluid intelligence, and cognitive function, *β* = −0.17, *SE* = 0.07, *t*(110) = −2.53, *p* = 0.013. We note that a similar numerical pattern was observed in the narrative recall analyses, suggesting that these results reflect a quantitative but not a qualitative shift in the brain-behavior relationship.

### Connectome based predictive modeling

In a planned exploratory analysis, we ran a CBPM analysis to see if we could predict narrative recall using the entire intrinsic functional connectome. The grand mean functional connectome for our sample is reported in the Supplemental Materials. The first analysis—where we used the entire 406 x 406 intrinsic functional connectome to predict narrative recall—resulted in a significant correlation between observed and predicted narrative recall scores (*r*_{observed, predicted}_ = 0.164, *p* = 0.027) – suggesting that there is a multivariate pattern within the broader functional connectome that is predictive of individual differences in narrative recall.

To ensure the robustness of this result, we first reran the CBPM analysis while controlling for nuisance variables when selecting connections to be used for predicting left out subjects (using the MATLAB function *partialcorr*, see Shen et al. (2017)). Specifically, we saw that the predictive performance of the intrinsic connectome held when controlling for age, self-reported sex, and average framewise displacement (*r*_{observed, predicted}_ = 0.1498, *p* < 0.01). All subsequent CBPM analyses were performed controlling for age, self-reported sex, and average framewise displacement. We next tested whether this result held when selecting different connection selection thresholds. In line with previous reports that the CBPM method is robust to connection selection threshold (Finn et al., 2015; Jangraw et al., 2018; Shen et al., 2017), rerunning the CBPM analysis using connection selection thresholds of *p* = [0.001, 0.005, 0.01, 0.5, 0.1] had similar outcomes (see Supplemental Materials). Finally, we examined whether these results were affected by the inclusion of global signal regression in our processing pipeline (see Supplemental Materials). Removing the global signal regression resulted in non-significant prediction performance, *r*_{observed, predicted}_ = 0.081, *p* = 0.099. Together, these results gave us confidence that our result was robust to most analytic decisions, with the exception that the inclusion of global signal regression appeared to increase predictive performance.

We then examined how combining functional connectomes from different tasks influenced our model. Previous research suggests that functional connectivity calculated by collapsing across resting-state and task scans increases the reliability and thus the predictive utility of individuals’ functional connectomes (Elliott et al., 2019). To confirm that this was the case in our data, we reran the CBPM analysis on functional connectomes calculated using the movie watching, resting-state, and sensorimotor task scans separately. Models built using functional connectomes from each task individually underperformed compared to the model built using the combined intrinsic functional connectome. Among the three tasks, the model built using functional connectomes from the movie-watching scan was the only one able to predict narrative recall better than chance. Interestingly, models built using resting-state data performed particularly poorly, with almost no relationship between observed and predicted narrative recall scores in out-of-sample data (see Supplemental Materials). These results support previous reports that suggested that functional connectivity estimates calculated across task and resting-state scans outperform those from short resting-state scans alone (Elliott et al., 2019).

To identify which networks were driving model performance, we ran a computational lesion analysis. In the computational lesion analysis, we reran our CBPM analysis excluding each network from the analysis in turn. Network importance in this computational lesion analysis was determined via a significant drop in model performance with the exclusion of a network and its connections. The results of our computational lesion analyses are reported in Table 2.

**Table 2:**
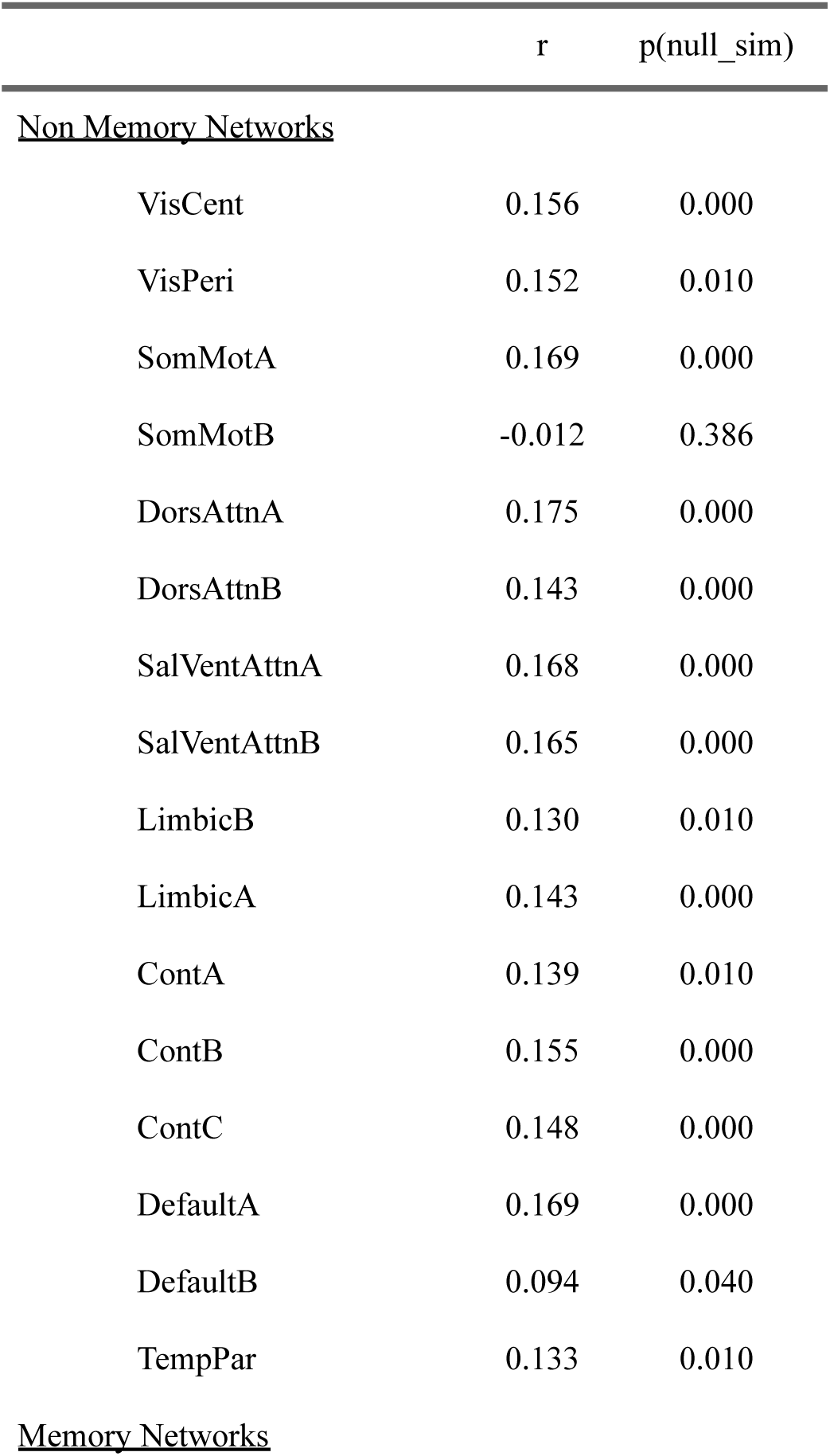
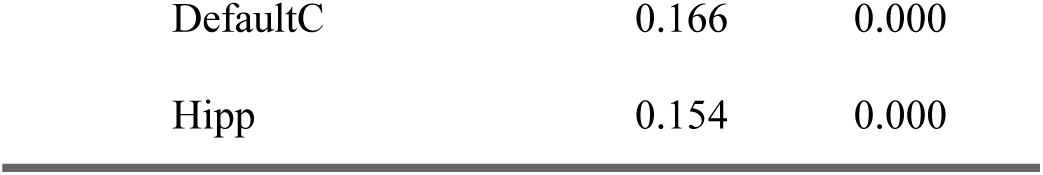
Computational Lesion Analysis. Excluding networks of regions traditionally related to episodic memory ability – i.e., the Default Mode Network C and the Hippocampus – have very little impact on model performance. Among non memory networks, only excluding Somatomotor Network B regions from the analysis resulted in a significant drop in model performance. r = Pearson’s correlation between observed and predicted narrative recall scores, p = proportion of null simulations that were more extreme than observed. See Yeo et al. (2011) and Schaefer et al. (2018) for descriptions of each network.

|  | r | p(null_sim) |
| --- | --- | --- |
| <u>Non Memory Networks</u> |  |  |
| VisCent | 0.156 | 0.000 |
| VisPeri | 0.152 | 0.010 |
| SomMotA | 0.169 | 0.000 |
| SomMotB | -0.012 | 0.386 |
| DorsAttnA | 0.175 | 0.000 |
| DorsAttnB | 0.143 | 0.000 |
| SalVentAttnA | 0.168 | 0.000 |
| SalVentAttnB | 0.165 | 0.000 |
| LimbicB | 0.130 | 0.010 |
| LimbicA | 0.143 | 0.000 |
| ContA | 0.139 | 0.010 |
| ContB | 0.155 | 0.000 |
| ContC | 0.148 | 0.000 |
| DefaultA | 0.169 | 0.000 |
| DefaultB | 0.094 | 0.040 |
| TempPar | 0.133 | 0.010 |
| Memory Networks |  |  |
| DefaultC | 0.166 | 0.000 |
| Hipp | 0.154 | 0.000 |

Exclusion of typical memory-related brain regions (i.e., DMN-C regions and the hippocampus) had little impact on model performance. Among non-memory related networks, the Somatomotor Network B was the only network whose exclusion led to a significant drop in model performance. To visualize the specific connections related to narrative recall, Figure 3 depicts the partial correlation between intrinsic functional connectivity and narrative recall scores for every connection in the functional connectome, averaged within each network. We also calculated the number and proportion of connections between each of our networks of interest that exceeded our connection selection threshold of *p* < 0.01 (see Supplemental Materials), as these constitute the connections that actually went into the CBPM analysis. These visualizations reveal a pattern of increases and decreases throughout the connectome associated with individual differences in narrative recall. Taken all together, these results suggest that memory ability related information was largely contained within connections of non-memory related brain regions

**Figure 3:**
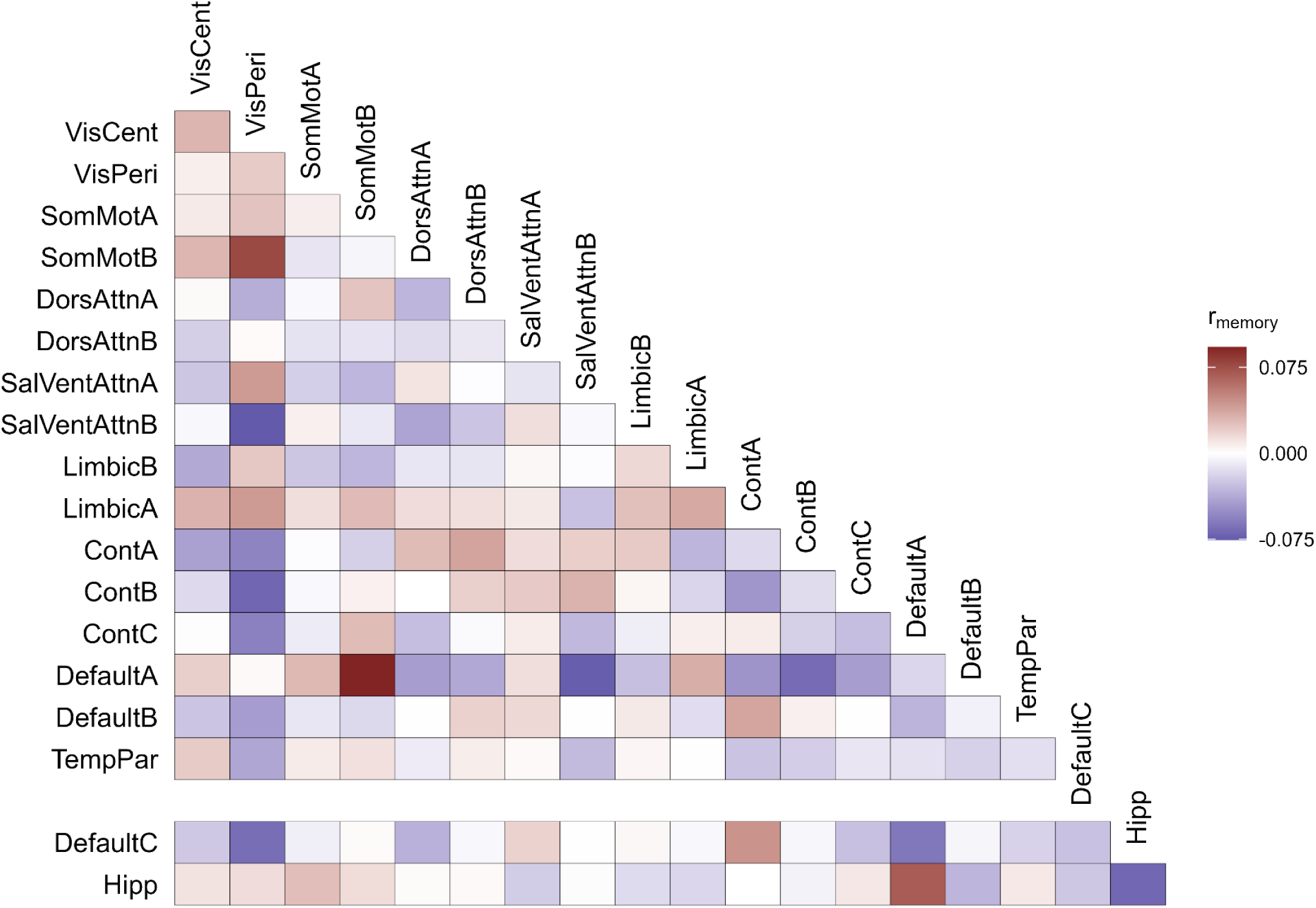
Evaluating Feature Importance. The partial correlation of connection strength with narrative recall scores after controlling for age, sex, and average in-scanner motion. DMN-C and hippocampal connections are offset to highlight these memory-related regions. See Yeo et al. (2011) and Schaefer et al. (2018) for descriptions of each network.

## 4 Discussion

The current report examined how individual differences in episodic memory ability are related to intrinsic functional connectivity, focusing specifically on regions in the default mode network (DMN-C) and the hippocampus that have been strongly implicated in episodic memory (Ritchey & Cooper, 2020). Across a sample of 243 individuals, we found little evidence that the strength of DMN-C and hippocampal functional connections was related to narrative recall performance. We did find evidence, however, that network-wide patterns of functional connectivity could predict individual differences in narrative recall. These predictive models were robust to analytic decisions and remained significant after controlling for age, self-reported sex, and average in-scanner movement. Probing these predictive models further revealed that superior narrative recall performance was related to functional connectivity of brain regions not typically related to memory ability – specifically, to somatomotor regions. Together, these results suggest that there is limited evidence connecting individual differences in memory to measures of intrinsic functional connectivity among regions typically associated with memory function.

The null results reported here were contrary to our a-priori hypotheses, and they contrast with previous reports linking changes in intrinsic functional connectivity of default mode network (Sneve et al., 2017; van Buuren et al., 2019) and hippocampal regions (Setton, Mwilambwe-Tshilobo, et al., 2022; Touroutoglou et al., 2015; L. Wang, LaViolette, et al., 2010; L. Wang, Negreira, et al., 2010) to episodic memory ability. One possible explanation for this set of results is that the strength of intrinsic functional connections might not be reliably related to episodic memory ability, despite its relationship to other measures of cognitive performance. Measures of intrinsic functional connectivity, often obtained from resting-state scans that do not include an explicit cognitive task, have been widely used to study individual differences in cognition. Strong evidence has accumulated over the years to suggest that these measures are related to behavioral phenotypes. Studies of the resting-state have found that there is a normative pattern of functional connections in the brain, such that brain regions form stable networks (between 7-17, Yeo et al., 2011). Recent work suggests that the majority of variability in the strength of these connections is attributable to stable individual differences away from this group-level pattern (as opposed to variation attributable to cognitive task or day-to-day variation; (Gratton et al., 2018). Furthermore, the strength of intrinsic connections has been shown to be predictive of a number of different behavioral phenotypes including neuroticism and extraversion (Hsu et al., 2018), trait-level anxiety (Z. Wang et al., 2021), fluid intelligence (Finn et al., 2015), creativity (Beaty et al., 2018), sustained attention (Rosenberg et al., 2016), and working memory ability (Avery et al., 2020). Patterns within the intrinsic functional connectome are so identifiable that they can be used to identify an individual from a group, acting as a sort of “brain fingerprint” (Finn et al., 2015). Thus, it seemed reasonable to hypothesize that individual differences in intrinsic functional connectivity would be related to memory ability.

Other work critiques this line of research, however, suggesting that “intrinsic” functional connectivity calculated in the resting-state is less useful for predicting individual differences compared with “active” functional connectivity calculated while participants complete cognitive tasks (Greene et al., 2018, 2020; Lin et al., 2021), especially tasks that are relevant to the predicted cognitive ability (Zhao et al., 2023). An analogy would be examining two cars and trying to determine which one is the better race car — you may not be able to tell the difference between an expensive race car and a Toyota Civic by looking under the hood when they are sitting in a garage, but you may be able to tell the difference when you examine how their mechanical parts perform when engaged in a race. The results of one of our supplemental analyses support this idea, with out-of-sample performance being significantly worse using the resting-state scan compared with the movie-watching or sensorimotor task data (see Supplemental Materials). A closely related critique also questions the use of resting state networks for understanding how the brain supports cognition. This critique specifically puts forth the idea that the most important unit of analysis when relating brain measures to cognition is not networks identified via low-frequency coactivation during rest, but networks identified via high frequency, transient coupling while participants complete highly controlled cognitive tasks (Cabeza & Moscovitch, 2013; Davis et al., 2017; Moscovitch et al., 2016). These “process specific alliances” often straddle resting-state network definitions, and in contrast to resting-state networks, they are thought to be able to flexibly assemble and disassemble depending on task demands. For example, interactions between the hippocampus and the left ventrolateral prefrontal cortex are thought to support successful episodic encoding (e.g., Wing et al., 2013), despite the fact that hippocampus is typically a member of the default mode network and left ventrolateral prefrontal cortex is not (Davis et al., 2017). Thus, it could be the case that individual differences in behavior manifest themselves in individual differences in the strength of these high frequency and cognition specific alliances and not in strength of low-frequency resting state networks. Here, we combined data from resting-state and task scans in order to estimate a “generalized” measure of functional connectivity (Elliott et al., 2019), which is thought to be a more reliable measure than those derived from short resting-state scans alone, but will necessarily obscure task-specific variation. When considering the scans separately, we found largely similar results across the different types of scans. However, there was evidence for significant connectome-based prediction only for the movie-watching scan, which is notable because processing event narratives in the context of movie-watching may be more relevant to the kinds of processes that support performance on narrative recall tasks (Jääskeläinen et al., 2021). Thus, although we did not find a relationship between memory ability and intrinsic measures of functional connectivity in memory related brain regions, we hypothesize that this relationship may be apparent when measuring brain activity while participants complete a memory-related task, or when incorporating brain measures that isolate a particular pattern of activity related to memory function (e.g., Richter et al., 2023).

In our data-driven analyses, we found that a multivariate brain-wide pattern of functional connectivity could predict individual differences in narrative recall reliably out-of-sample. Our predictive analyses suggest that the brains of superior rememberers are characterized, in part, by decoupling of somatomotor regions and DMN-A regions and increased communication between somatomotor regions and visual processing regions. We speculate that the involvement of somatomotor regions in predicting memory ability might be explained by variation in goal-directed attention in the sensorimotor task, as successful attention in this task might be related to attentional factors contributing to individual differences in memory (Unsworth, 2019). The decoupling of default and sensory regions is in line with previous results that likewise showed evidence the superior rememberers were characterized by decoupling between default mode and sensory networks (Sneve et al., 2017). Other studies, however, have shown that general increases in task-related DMN connectivity are related to episodic memory ability (King et al., 2015) and that lower MTL-DMN and higher DMN-frontoparietal control network connections are related to memory ability (van Buuren et al., 2019). A recent, well-powered study identified multivariate patterns of associations between individual differences in autobiographical memory and functional connectivity among DMN regions, including the temporal pole and hippocampus, although these relationships were largely outside of the neocortical regions most commonly associated with episodic memory (Setton, Mwilambwe-Tshilobo, et al., 2022). What is clear from the results presented here and the literature to date is that information about memory ability in healthy young adults is likely contained within the functional connectome beyond brain networks classically linked to episodic remembering (Lin et al., 2021). The exact pattern of whole brain connectivity that predicts memory ability, however, remains inconsistent from study to study. As such, we refrain from drawing strong conclusions about the specific connections related to memory in the present study.

An important factor that may have influenced our results is our choice of memory measure. Here, we used a standard neuropsychological measure of memory that assessed an individual’s ability to recall the details of a short narrative – the logical memory subtest of the Wechsler Memory Scale (Weschler, 1999). The logical memory subtest of the Wechsler Memory Scale was chosen because it was available for the largest number of subjects in the Cam-CAN dataset, compared to other included memory measures, and this measure has been previously shown to correlate with individual differences in brain activity in response to event boundaries during the Cam-CAN movie watching scan (Reagh et al., 2020). Narrative recall has also been linked specifically to the function of the DMN, such that there is increased activity and functional integration among DMN regions during recall of event narratives (Lee et al., 2020; Ritchey & Cooper, 2020). Reliance on a single memory measure to capture an individual’s memory ability, however, may be problematic given previous research suggesting that individuals have aptitudes for different types of memory tasks (Unsworth, 2019). We speculate that differences in how memory ability is operationalized could explain the mixed state of the literature – all of the previous research relating individual differences in memory ability and functional connectivity have taken a unique approach to operationalizing memory ability (King et al., 2015; Lin et al., 2021; Sneve et al., 2017; van Buuren et al., 2019). King and colleagues (2015) and Sneve and Colleagues (2017) relied on performance in source memory recognition tasks; Lin and colleagues (2021) relied on performance on a remember/know/new recognition memory paradigm; van Burren and colleagues (2019) relied on performance on an object-location memory task; Setton and colleagues (2022) used the number of internal details generated during an autobiographical memory interview. The brain areas supporting retrieval depend in part on the content, modality, and format of memory tasks (Diana et al., 2007), consistent with the idea that memory performance is supported by a combination of task-specific and task-general processes (Rugg & Vilberg, 2013). Such differences in the content and format of memory tasks may explain why there has not been one consistent neural signature associated with performance. Here, we attempted to test the generalizability of our results by creating a composite measure of memory ability in a subset of our sample, again finding little evidence for a relationship between DMN functional connectivity and memory ability. However, this analysis incorporated only one additional memory measure and was available for only about half of our sample. Future research on individual differences in memory ability could use multiple memory measures to get an unbiased measure of individuals’ overall memory ability, or systematically compare memory measures that are likely to correspond with dissociable neural processes (e.g., Ngo et al., 2021).

While these previous studies have considered individual differences in objective evaluations of participants’ memory, other studies have looked at measures of individuals’ subjective evaluations of their own memory when relating individual differences to brain function (Petrican et al., 2020; Sheldon et al., 2016). Sheldon and colleagues (2016), for example, examined the relationship between the strength of functional connectivity of the parahippocampal cortex and subjective reports of episodic memory tendencies measured by the Survey of Autobiographical Memory (SAM) survey (Palombo et al., 2013). The SAM measures participants’ self-reported mnemonic traits, measuring their tendency, for example, to remember specific event and contextual details when recalling events (episodic subscale) versus their tendency to remember facts about oneself, events, or the world that lack contextual detail (semantic subscale). There is some evidence that individual differences in self-reported mnemonic traits measured by the episodic subscale of the SAM are associated with a similar resting-state functional connectivity profile as individual differences measured using a visual, laboratory based episodic memory task (Petrican et al., 2020). Future work should consider the relationship between subjective and objective measures of memory function (Clark & Maguire, 2020; e.g., Cooper & Ritchey, 2022; Setton, Lockrow, et al., 2022) in the context of how these measures relate to individual differences in brain function.

An intriguing future direction is to look at other facets of brain connectivity and how they may be related to episodic memory ability. There are at least three facets of brain connectivity that could theoretically support individual differences in cognition: variability in connectional strength, variability in the spatial localization of brain regions, and variability in large-scale network topology (i.e., large scale networks have different sets of constituent nodes across subjects (Gordon & Nelson, 2021)). The current study, like many of the studies that have come before, focused on how variability of the strength of functional connections related to individual differences in memory ability. Recent work, however, suggests that there are substantial individual differences in the size and topological organization of functional brain areas (Gordon, Laumann, Adeyemo, et al., 2017; Gordon & Nelson, 2021; Kong et al., 2019; Laumann et al., 2015). Gordon and Nelson (2021) display a striking example of this type of idiosyncrasy, where the posterior medial precuneus node of the default mode network for one subject is translated along the cortical surface such that the node wraps around to the lateral side of the brain. Recent work has also identified individual differences in large-scale network topology, such that nodes that are the same across individuals are a part of different large scale networks (Gordon & Nelson, 2021; Laumann et al., 2015; Seitzman et al., 2019). Such “network variants” appear to occur in specific regions of the brain, particularly in default mode regions, and are observed in some datasets in approximately 33% of individuals (Seitzman et al., 2019). These differences in the spatial topography of functional nodes on the cortical surface also appear to have behavioral relevance. Individuals that have similar spatial network topographies also perform similarly on cognitive tasks, so much so that individual behavior can be predicted out of sample using similarity in brain wide network topography and network size (Kong et al. 2019). An intriguing possibility is that individual differences in memory ability could be related to the presence or absence of network variants in default mode regions.

In conclusion, we found little evidence in support of a relationship between individual differences in narrative recall and intrinsic functional connectivity among cortical and hippocampal networks commonly associated with episodic memory. We did find a multivariate brain wide pattern of functional connectivity that was predictive of narrative recall, characterized by decoupling of somatomotor regions from default mode regions. Our findings agree with previous research that suggests that information about memory ability is contained in functional connections in regions outside of regions classically linked to episodic memory. The exact nature of this brain connectivity pattern that predicts memory ability remains unclear. We believe this is due to variability in how memory ability is operationalized and the low power of previous studies both in terms of number of subjects collected and in the amount of data collected per subject. Future research on the relationship between individual differences in memory and functional brain networks should incorporate multiple measures to operationalize memory ability unbiased toward a particular task (Unsworth, 2019), collect a large amount of data (Gordon, Laumann, Adeyemo, et al., 2017; Marek et al., 2022), especially task-related data (Greene et al., 2018), and examine additional facets of brain organization that may better capture variability across individuals (Gordon & Nelson, 2021).

## Data and Code Availability

The current study reports data that is openly available from the Cambridge Center for Aging Neuroscience (Shafto et al., 2014; Taylor et al., 2017). Custom code used to process and analyze these data is available in the following repository: https://github.com/memobc/paper-CamCanIDs

## Author Contributions

K.K.: conceptualization, formal analysis, software, visualization, writing-original draft, writing-review & editing. M.R.: conceptualization, funding acquisition, supervision, writing-review & editing.

## Acknowledgements

This work was supported by NIH grant R01MH125990 and NSF grant BCS-2047415 awarded to M.R. We would like to acknowledge Tingwei Hu for assisting with quality assurance checks on the MRI data included in this report. We additionally thank Elizabeth Kensinger, Ehri Ryu, and Chris Bird for providing feedback on previous versions of this manuscript, which were included as part of the first author’s dissertation.

## Supplemental Materials

**Figure S1:**
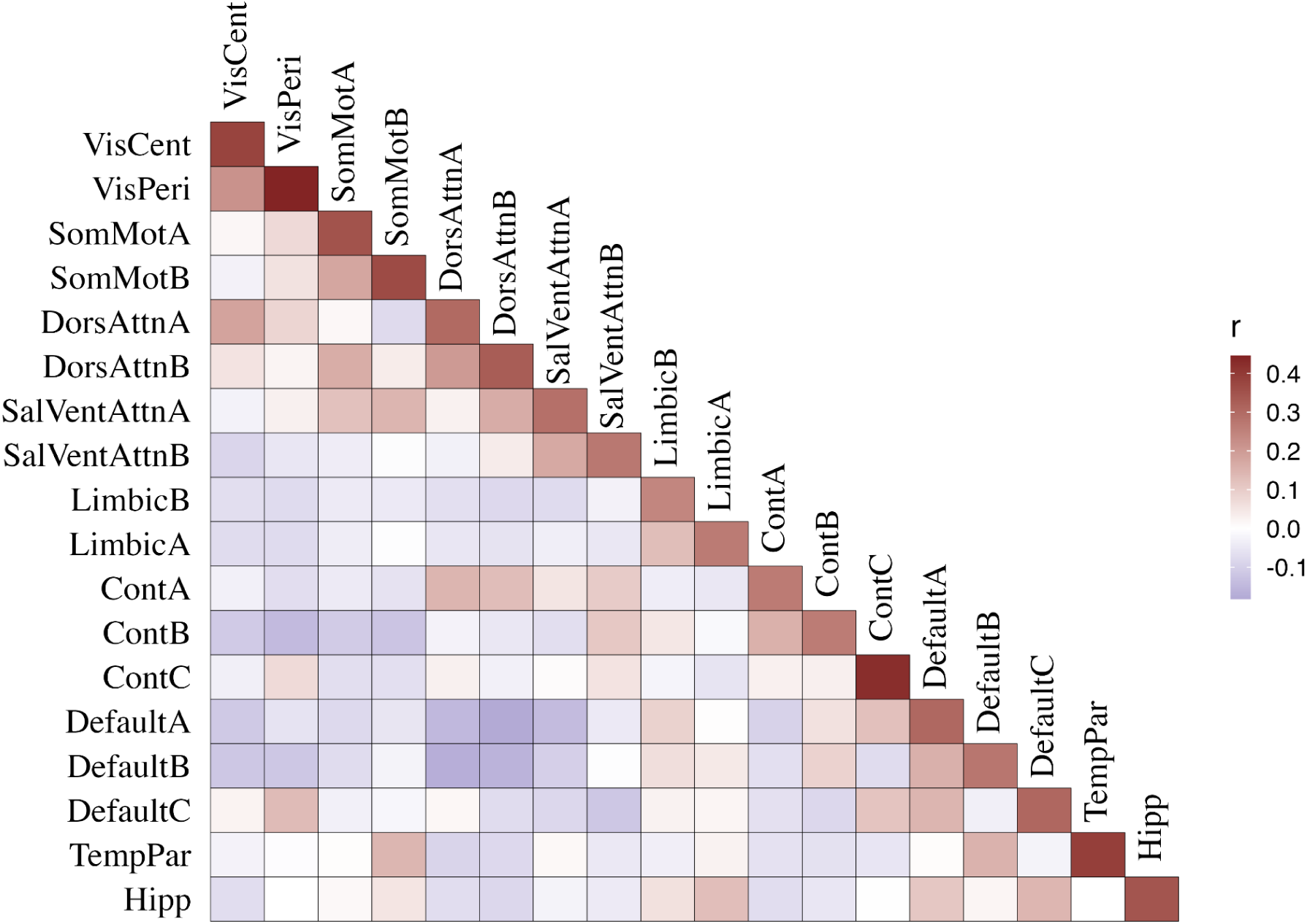
Grand mean connectivity matrix. Here we report the grand mean functional connectivity matrix, calculated by averaging functional connectivity estimates between our networks of interest across all subjects included in our analysis. As expected, regions included in the same functional network display increased functional connectivity. See Yeo et al. (2011) and Schaefer et al. (2018) for descriptions of each network.

**Figure S2:**
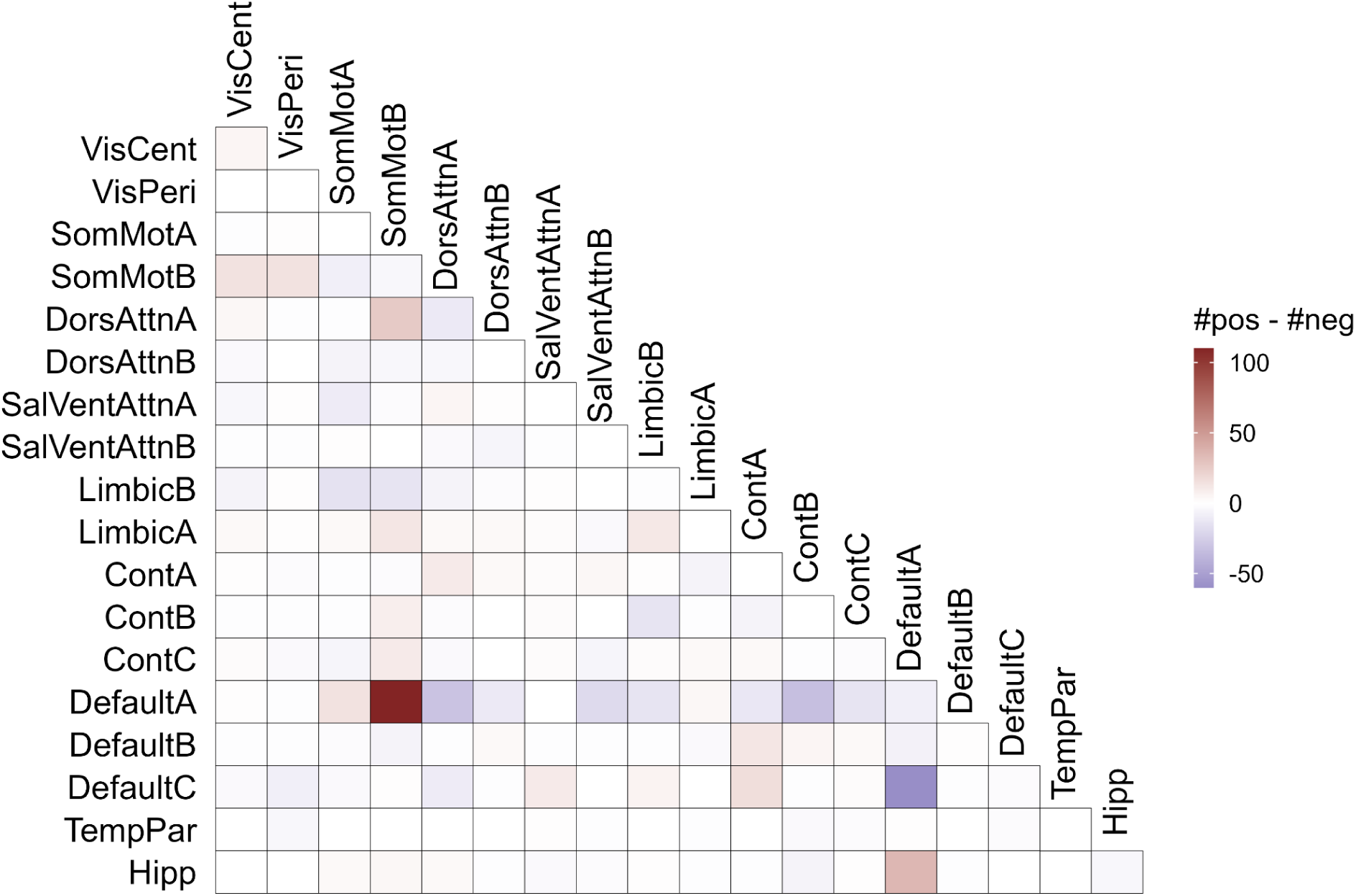
Number of positive - negative connections that entered our predictive model. For each pair of our networks of interest, we calculated the number of connections that positively correlated with memory ability at a *p* < 0.01 (our connection selection threshold, see text) and subtracted the number of connections that negatively correlated with memory ability at a *p* < 0.01.

**Figure S3:**
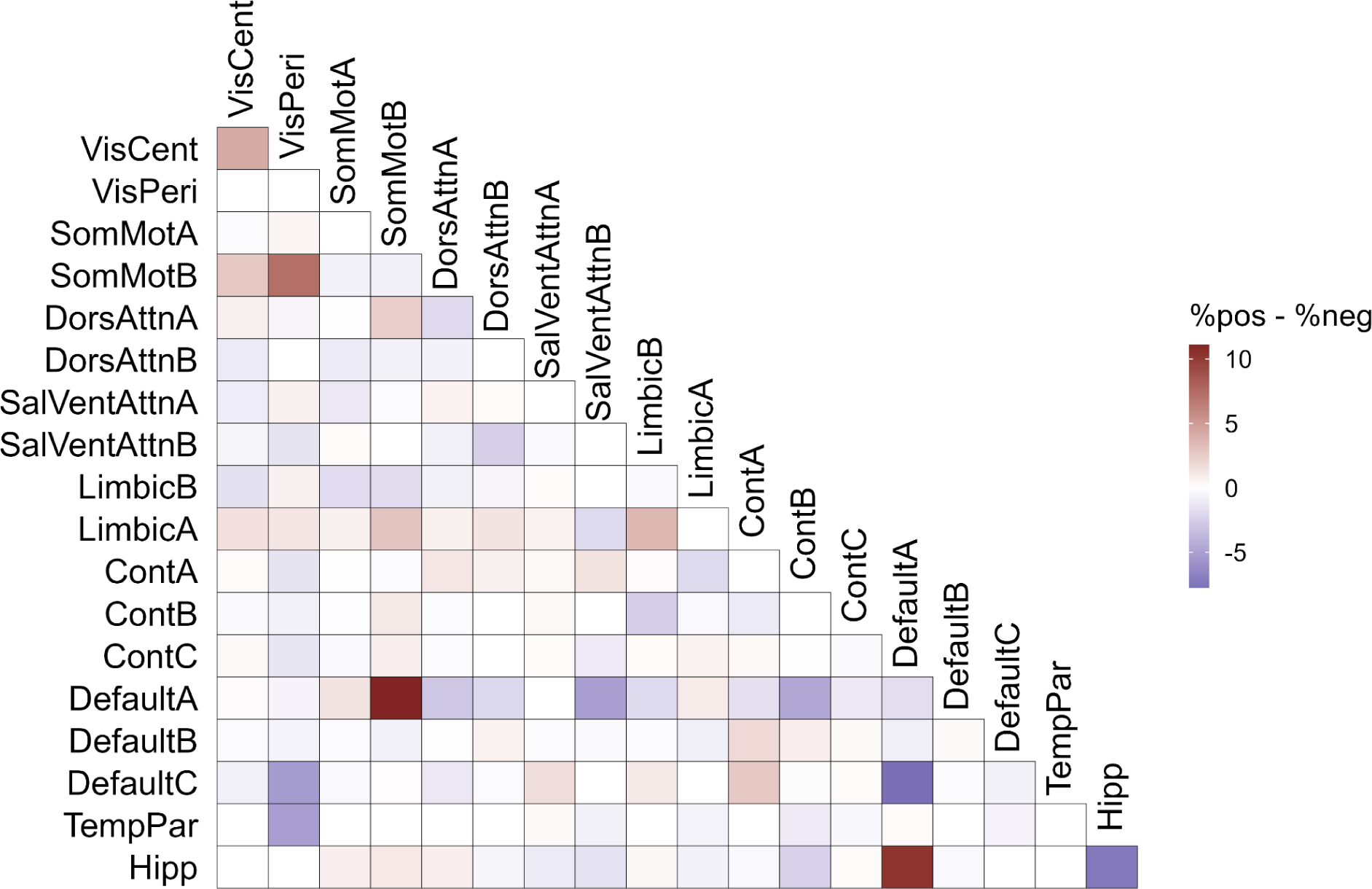
Proportion of connections that entered our predictive model. For each pair of our networks of interest, we calculated the proportion of connections (of all possible connections involving that pair) that positively correlated with memory ability at a *p* < 0.01 (our connection selection threshold, see text) and subtracted the proportion of connections that negatively correlated with memory ability at a *p* < 0.01.

### Analyses excluding global signal regression

Global signal regression (GSR) involves statistically removing the global mean signal from the timecourse of each voxel in the brain prior to calculating functional connectivity between brain regions (see Murphy & Fox, 2017 for a review). GSR was originally thought to be an effective way to remove artifactual signals (e.g., motion) from voxel time series and was implemented in task-based fMRI experiments. An important caveat to this procedure is that it mathematically mandates negative connectivity estimates between brain regions (Murphy et al., 2009), muddying the interpretation of resulting anti-correlations. Recent research, however, suggests that GSR helps analyses whose aim is to predict behavior using functional connectivity estimates (Finn & Bandettini, 2021; Li et al., 2019), likely related to its efficacy in controlling head-related motion artifacts and other noise sources affecting the global signal (Satterthwaite et al., 2019). In our pre-registration, we did not include GSR in our preprocessing pipeline, though we have since been convinced by these recent studies that this step may be important for individual difference analyses. In the main text, we report results including GSR, and for completeness, we reran our analyses *excluding* GSR from our preprocessing pipeline and report the results below. Removing GSR had no impact on our pre-registered, hypothesis-driven analyses targeting the DMN-C (see Supplemental Table 1-3). However, there were two differences in the other results: First, removing GSR results in a statistically significant relationship between average hippocampal connectivity on memory ability, wherein average hippocampal connectivity was inversely related to memory ability (see Supplemental Table 4). Second, removing GSR resulted in a substantially weaker ability to predict memory ability in the CBPM analysis (*r*_{observed, predicted}_ = 0.081, *p* = 0.099), controlling for age, sex, and framewise displacement. In other words, it appears that including GSR reduced the relationship between hippocampal connectivity and memory, but it improved predictive performance in the overall CBPM analysis.

One possible explanation for this discrepancy is that, while GSR effectively minimizes the influence of head motion on functional connectivity estimates, it does so in a distance-dependent way (Satterthwaite et al., 2013), controlling for motion-related covariation among long-range connections more effectively than among short-range connections (Satterthwaite et al., 2019). We speculate without GSR, unaccounted motion artifacts, especially among long-range connections, might have resulted in a spurious connection between hippocampal connectivity and memory ability, while also interfering with the generalizability of predictions in the cross-validated CBPM analysis. Alternatively, there may be an actual trend toward a negative relationship between hippocampal connectivity and memory ability, as we also observed this relationship when using a composite score of memory ability (see Results) with GSR. This pattern would be unexpected, based on past research showing that hippocampal connectivity is positively related to memory ability (Setton, Mwilambwe-Tshilobo, et al., 2022; Touroutoglou et al., 2015; L. Wang, LaViolette, et al., 2010; L. Wang, Negreira, et al., 2010). However, in one study using a relatively small sample, there was evidence for a negative relationship between memory detail and hippocampal functional connectivity, specifically with the medial prefrontal cortex, but only among older adults (Matijevic et al., 2022). While it is important to consider the full spectrum of possible results here, due to the unexpectedness of this result and its sensitivity to different analysis decisions, we think that it is best to remain cautious in interpreting this result.

**Table S1:** Regression results of average within DMN-C connectivity on memory ability removing GSR from our analysis pipeline. within = average strength of connection among DMN-C regions; acer = cognitive function score, cattell = fluid intelligence score. ^1^*p<0.05; **p<0.01; ***p<0.001. ^2^SE = Standard Error. ^3^Female = 0, Male = 1.

|  | Model 1 |  | Model 2 |  | Model 3 |  |
| --- | --- | --- | --- | --- | --- | --- |
| Characteristic | Beta<sup>1</sup> | SE<sup>2</sup> | Beta<sup>1</sup> | SE<sup>2</sup> | Beta<sup>1</sup> | SE<sup>2</sup> |
| within | -0.29 | 0.225 | -0.27 | 0.225 | -0.18 | 0.204 |
| age |  |  | -0.05 | 0.027 | -0.05 | 0.026 |
| sex<sup>3</sup> |  |  | -0.91* | 0.446 | -1.00* | 0.414 |
| fd |  |  | -6.36 | 5.70 | 1.98 | 5.21 |
| acer |  |  |  |  | 1.10*** | 0.243 |
| cattell |  |  |  |  | 0.83*** | 0.237 |
| No. Obs. | 243 |  | 243 |  | 235 |  |
| R<sup>2</sup> | 0.007 |  | 0.048 |  | 0.238 |  |

**Table S2:** Regression results of average DMN-C--DMN-A connectivity on memory ability removing GSR from our analysis pipeline. between = average strength of connection between DMN-C and DMN-A regions; acer = cognitive function score, cattell = fluid intelligence score. ^1^*p<0.05; **p<0.01; ***p<0.001. ^2^SE = Standard Error. ^3^Female = 0, Male = 1.

|  | Model 4 |  | Model 5 |  | Model 6 |  |
| --- | --- | --- | --- | --- | --- | --- |
| Characteristic | Beta<sup>1</sup> | SE<sup>2</sup> | Beta<sup>1</sup> | SE<sup>2</sup> | Beta<sup>1</sup> | SE<sup>2</sup> |
| between | -0.35 | 0.224 | -0.22 | 0.231 | -0.07 | 0.208 |
| age |  |  | -0.04 | 0.027 | -0.05 | 0.026 |
| sex<sup>3</sup> |  |  | -0.95* | 0.444 | -1.02* | 0.414 |
| fd |  |  | -5.62 | 5.86 | 1.93 | 5.33 |
| acer |  |  |  |  | 1.11*** | 0.244 |
| cattell |  |  |  |  | 0.82*** | 0.237 |
| No. Obs. | 243 |  | 243 |  | 235 |  |
| R<sup>2</sup> | 0.010 |  | 0.046 |  | 0.235 |  |

**Table S3:**
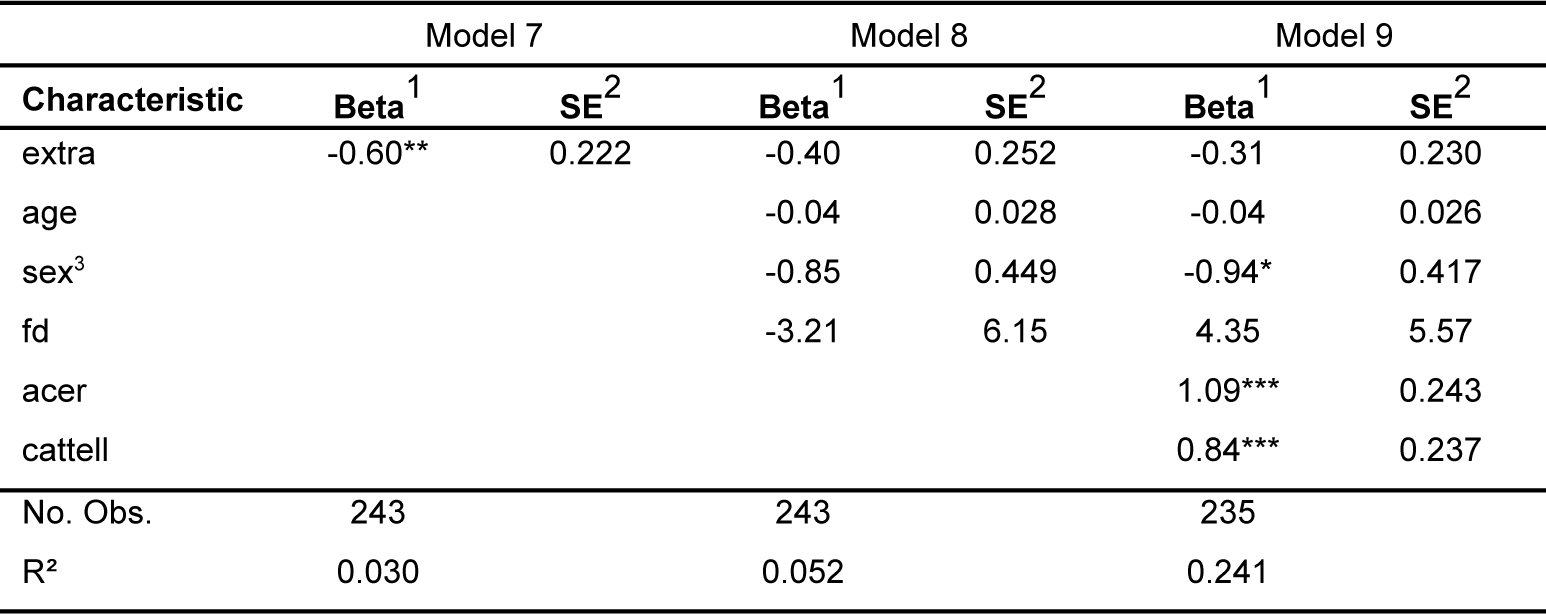
Regression results of average DMN-C connectivity with the rest of the brain on memory ability removing GSR from our analysis pipeline. extra = average strength of connection between DMN-C regions and regions not in the DMN-C or DMN-A; acer = cognitive function score, cattell = fluid intelligence score. ^1^*p<0.05; **p<0.01; ***p<0.001. ^2^SE = Standard Error. ^3^Female = 0, Male = 1.

**Table S4:**
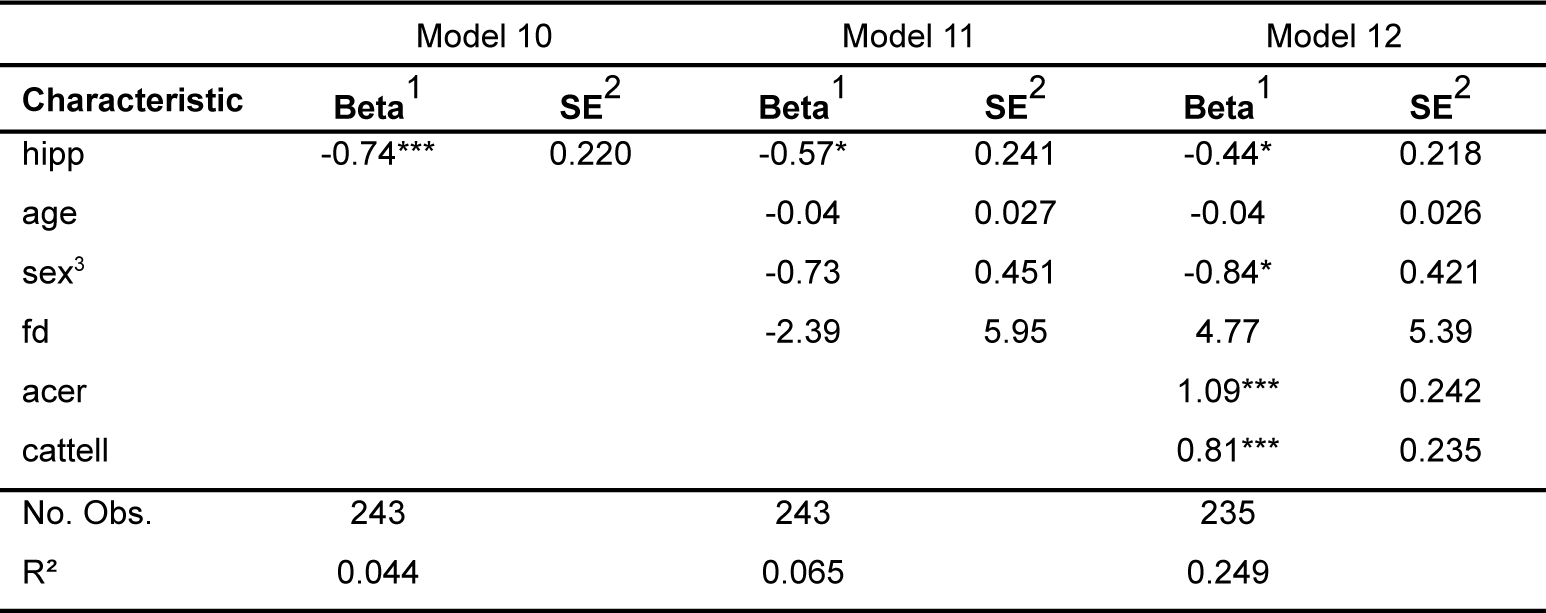
Regression results of average hippocampal connectivity on memory ability removing GSR from our analysis pipeline. hipp = average strength of connection of hippocampal regions; acer = cognitive function score, cattell = fluid intelligence score. ^1^*p<0.05; **p<0.01; ***p<0.001. ^2^SE = Standard Error. ^3^Female = 0, Male = 1.

|  | Model 10 |  | Model 11 |  | Model 12 |  |
| --- | --- | --- | --- | --- | --- | --- |
| Characteristic | Beta<sup>1</sup> | SE<sup>2</sup> | Beta<sup>1</sup> | SE<sup>2</sup> | Beta<sup>1</sup> | SE<sup>2</sup> |
| hipp | -0.74*** | 0.220 | -0.57* | 0.241 | -0.44* | 0.218 |
| age |  |  | -0.04 | 0.027 | -0.04 | 0.026 |
| sex<sup>3</sup> |  |  | -0.73 | 0.451 | -0.84* | 0.421 |
| fd |  |  | -2.39 | 5.95 | 4.77 | 5.39 |
| acer |  |  |  |  | 1.09*** | 0.242 |
| cattell |  |  |  |  | 0.81*** | 0.235 |
| No. Obs. | 243 |  | 243 |  | 235 |  |
| R<sup>2</sup> | 0.044 |  | 0.065 |  | 0.249 |  |

### Impact of the connection selection threshold used in CBPM

Our study used Connectome Based Predictive Modeling (CBPM; Shen et al., 2017) to determine if information contained within the functional connectome is useful for predicting memory ability. This approach involves setting an arbitrary threshold for defining which connections are used to make behavioral predictions. Analyses reported in this manuscript used a connection selection threshold of *p* <= .01. It is unclear, however, whether this arbitrary choice of threshold has any impact on our results. We reran our key CBPM analysis (i.e., predicting memory ability while controlling for age, sex, and average framewise displacement) using a range of connection selection thresholds: *p* = [0.001 0.005 0.01 0.05 0.1]. Results are reported in Supplemental Figure 4. Analyses using all selection thresholds were statistically significant with one exception – when the selection threshold was set to *p* < 0.005. It is currently unclear why this specific analysis failed — selecting an even stricter threshold (i.e., *p* < 0.001) resulted in restored predictive performance. We take this pattern of results as evidence that the CBPM approach is robust to connection selection threshold, in line with previous reports (Finn et al., 2015; Jangraw et al., 2018; Shen et al., 2017).

**Figure S4:**
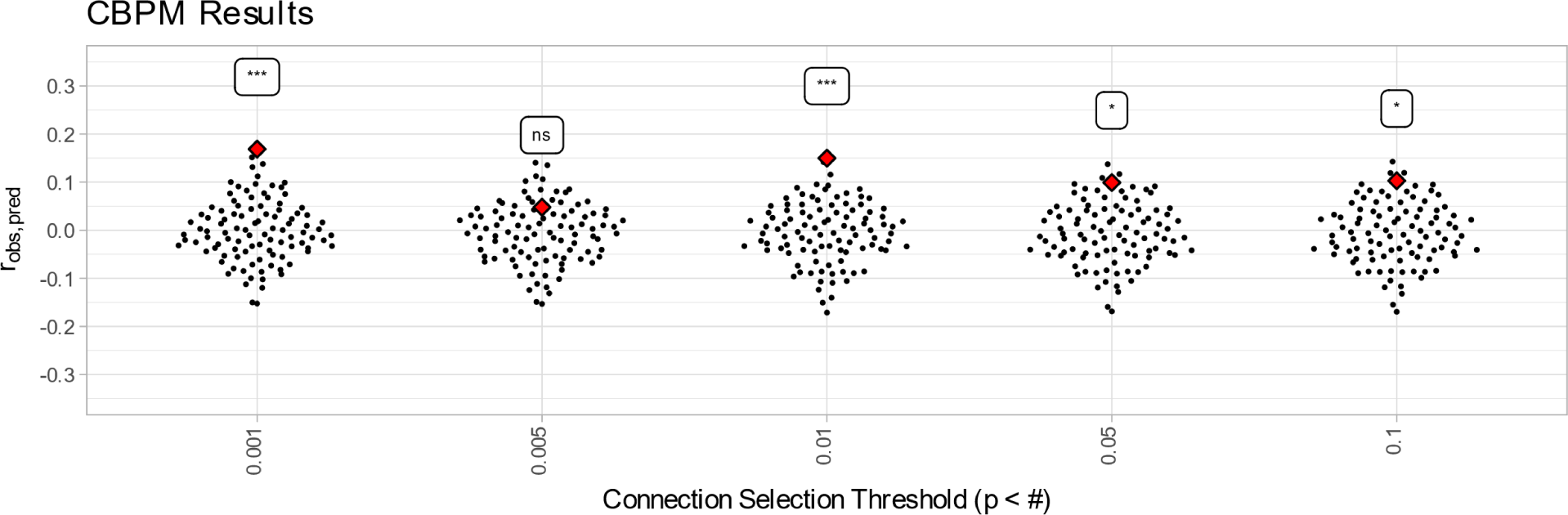
CBPM results are robust to selection of connection selection threshold in the CBPM analysis. Red diamonds indicate observed results, black dots indicate results of 100 null simulations. *** *p* <= 0.001, ** *p* <= 0.01, * *p* <= 0.05, ns = *p* > 0.05.

### Combining data across tasks appears to improve our ability to predict memory ability out of sample

Our analyses combined data across tasks to increase the reliability of our functional connectivity estimates (Elliott et al., 2019). This approach assumes that intrinsic functional connectivity would not vary substantially across tasks. Indeed, a recent study suggests that variability in the functional connectome is dominated by a normative pattern, patterns unique to individuals, and patterns unique to how individuals complete certain tasks (Gratton et al., 2018). To test the validity of this approach, we calculated the functional connectome separately for each task for each subject (i.e., movie watching, rest, sensorimotor) and correlated the resulting task-specific functional connectomes. Supplemental Table 5 displays the mean, standard deviation, minimum, and maximum similarity for each pair of tasks across our sample of 243 subjects. All correlations between tasks were performed on the subset of subjects that had a pair of valid scans. The movie-watching connectome was notably less similar to the rest and sensorimotor task connectomes. We suspect that this could be due to stimulus-driven changes in brain activation.

**Table S5:** How similar are connectomes calculated using data from different tasks? Similarity between connectomes was calculated using a Pearson’s correlation. N = number of subjects with valid scans for both scans in this pair, min = minimum Peasrson’s correlation between connectomes in task pair, max = maximum Pearson’s correlation between connectomes in task pair, mean = average Pearson’s correlation between connectomes in task pair.

| Task Pair | N | min | max | mean | sd |
| --- | --- | --- | --- | --- | --- |
| movie-smt | 240 | 0.20 | 0.69 | 0.51 | 0.06 |
| rest-movie | 227 | 0.18 | 0.66 | 0.49 | 0.06 |
| rest-smt | 226 | 0.47 | 0.79 | 0.63 | 0.07 |

To see how this impacted our results, we reran our CBPM analyses using functional connectomes calculated using only the movie-watching data (“movie”), only the resting-state data (“rest”), and only the sensorimotor (“smt”) tasks data. The results of these CBPM analyses are reported in Supplemental Figure 5. Functional connectomes calculated using the resting-state (*r*_{observed, predicted}_ = −0.007, *p* = 0.39) and sensorimotor task (*r*_{observed, predicted}_ = 0.083, *p* = 0.12) data were insufficient for predicting memory ability. Functional connectomes calculated using the movie watching data, however, were sufficient for predicting memory ability (*r*_{observed, predicted}_ = 0.113, *p* = 0.03). Interestingly, all the task specific predictive models performed worse compared with our predictive model that used a combined “intrinsic” connectome for each subject by averaging across tasks (*r*_{observed, predicted}_ = 0.1498, *p* < 0.01).

**Figure S5:**
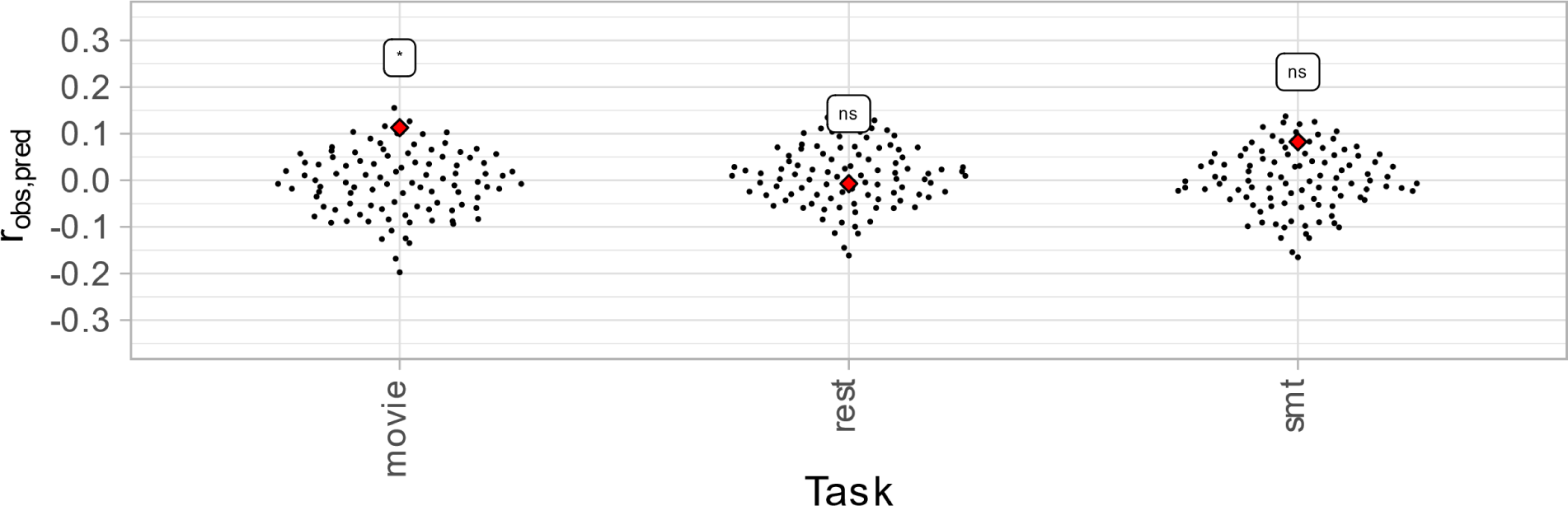
How do predictive models built using connectomes from individual tasks compare to models built using an “intrinsic” connectome, calculated by averaging across tasks? Movie = movie watching, rest = resting-state, smt = sensorimotor task.

### Detailed description of the fMRIPrep pipeline

The following is an edited version of the recommended boilerplate output by *fMRIPrep* after processing our data. The original boilerplate contained redundant descriptions of the operations performed by the software. What appears below is a detailed description of the processing steps with the redundant descriptions removed. Results included in this manuscript come from preprocessing performed using *fMRIPrep 20.2.0* (Esteban, Markiewicz, et al. (2018); Esteban, Blair, et al. (2018); RRID:SCR_016216), which is based on *Nipype 1.5.1* (Gorgolewski et al. (2011); Gorgolewski et al. (2018); RRID:SCR_002502).

### Anatomical data preprocessing

A total of 1 T1-weighted (T1w) images were found within the input BIDS dataset. The T1-weighted (T1w) image was corrected for intensity non-uniformity (INU) with N4BiasFieldCorrection (Tustison et al. 2010), distributed with ANTs 2.3.3 (Avants et al. 2008, RRID:SCR_004757), and used as T1w-reference throughout the workflow. The T1w-reference was then skull-stripped with a *Nipype* implementation of the antsBrainExtraction.sh workflow (from ANTs), using OASIS30ANTs as target template. Brain tissue segmentation of cerebrospinal fluid (CSF), white-matter (WM) and gray-matter (GM) was performed on the brain-extracted T1w using fast (FSL 5.0.9, RRID:SCR_002823, Zhang, Brady, and Smith 2001). Volume-based spatial normalization to one standard space (MNI152NLin2009cAsym) was performed through nonlinear registration with antsRegistration (ANTs 2.3.3), using brain-extracted versions of both T1w reference and the T1w template. The following template was selected for spatial normalization: *ICBM 152 Nonlinear Asymmetrical template version 2009c* [Fonov et al. (2009), RRID:SCR_008796; TemplateFlow ID: MNI152NLin2009cAsym].

### Functional data preprocessing

For each of the 3 BOLD runs found per subject (across all tasks and sessions), the following preprocessing was performed. First, a reference volume and its skull-stripped version were generated from the shortest echo of the BOLD run using a custom methodology of *fMRIPrep*. A B0-nonuniformity map (or *fieldmap*) was estimated based on a phase-difference map calculated with a dual-echo GRE (gradient-recall echo) sequence, processed with a custom workflow of *SDCFlows* inspired by the epidewarp.fsl script and further improvements in HCP Pipelines (Glasser et al. 2013). The *fieldmap* was then co-registered to the target EPI (echo-planar imaging) reference run and converted to a displacements field map (amenable to registration tools such as ANTs) with FSL’s fugue and other *SDCflows* tools. Based on the estimated susceptibility distortion, a corrected EPI (echo-planar imaging) reference was calculated for a more accurate co-registration with the anatomical reference. The BOLD reference was then co-registered to the T1w reference using flirt (FSL 5.0.9, Jenkinson and Smith 2001) with the boundary-based registration (Greve and Fischl 2009) cost-function.

Co-registration was configured with nine degrees of freedom to account for distortions remaining in the BOLD reference. Head-motion parameters with respect to the BOLD reference (transformation matrices, and six corresponding rotation and translation parameters) are estimated before any spatiotemporal filtering using mcflirt (FSL 5.0.9, Jenkinson et al. 2002). BOLD runs were slice-time corrected using 3dTshift from AFNI 20160207 (Cox and Hyde 1997, RRID:SCR_005927). The BOLD time-series (including slice-timing correction when applied) were resampled onto their original, native space by applying a single, composite transform to correct for head-motion and susceptibility distortions. These resampled BOLD time-series will be referred to as *preprocessed BOLD in original space*, or just *preprocessed BOLD*. A T2* map was estimated from the preprocessed BOLD by fitting to a monoexponential signal decay model with nonlinear regression, using T2*/S0 estimates from a log-linear regression fit as initial values. For each voxel, the maximal number of echoes with reliable signal in that voxel were used to fit the model. The calculated T2* map was then used to optimally combine preprocessed BOLD across echoes following the method described in (Posse et al. 1999). The optimally combined time series was carried forward as the *preprocessed BOLD*. The BOLD time-series were resampled into standard space, generating a *preprocessed BOLD run in MNI152NLin2009cAsym space*. Several confounding time-series were calculated based on the *preprocessed BOLD*: framewise displacement (FD), DVARS and three region-wise global signals. FD was computed using two formulations following Power (absolute sum of relative motions, Power et al. (2014)) and Jenkinson (relative root mean square displacement between affines, Jenkinson et al. (2002)). FD and DVARS are calculated for each functional run, both using their implementations in *Nipype* (following the definitions by Power et al. 2014). The three global signals are extracted within the CSF, the WM, and the whole-brain masks. Additionally, a set of physiological regressors were extracted to allow for component-based noise correction (*CompCor*, Behzadi et al. 2007). Principal components are estimated after high-pass filtering the preprocessed BOLD time-series (using a discrete cosine filter with 128s cut-off) for the two *CompCor* variants: temporal (tCompCor) and anatomical (aCompCor). tCompCor components are then calculated from the top 2% variable voxels within the brain mask. For aCompCor, three probabilistic masks (CSF, WM and combined CSF+WM) are generated in anatomical space. The implementation differs from that of Behzadi et al. in that instead of eroding the masks by 2 pixels on BOLD space, the aCompCor masks are subtracted a mask of pixels that likely contain a volume fraction of GM. This mask is obtained by thresholding the corresponding partial volume map at 0.05, and it ensures components are not extracted from voxels containing a minimal fraction of GM. Finally, these masks are resampled into BOLD space and binarized by thresholding at 0.99 (as in the original implementation). Components are also calculated separately within the WM and CSF masks. For each CompCor decomposition, the *k* components with the largest singular values are retained, such that the retained components’ time series are sufficient to explain 50 percent of variance across the nuisance mask (CSF, WM, combined, or temporal). The remaining components are dropped from consideration. The head-motion estimates calculated in the correction step were also placed within the corresponding confounds file. The confound time series derived from head motion estimates and global signals were expanded with the inclusion of temporal derivatives and quadratic terms for each (Satterthwaite et al. 2013). Frames that exceeded a threshold of 0.5 mm FD or 1.5 standardized DVARS were annotated as motion outliers. All resamplings can be performed with *a single interpolation step* by composing all the pertinent transformations (i.e. head-motion transform matrices, susceptibility distortion correction when available, and co-registrations to anatomical and output spaces). Gridded (volumetric) resamplings were performed using antsApplyTransforms (ANTs), configured with Lanczos interpolation to minimize the smoothing effects of other kernels (Lanczos 1964). Non-gridded (surface) resamplings were performed using mri_vol2surf (FreeSurfer).

Many internal operations of *fMRIPrep* use *Nilearn* 0.6.2 (Abraham et al. 2014, RRID:SCR_001362), mostly within the functional processing workflow. For more details of the pipeline, see the fMRIPrep documentation.

## Footnotes

1 Global signal regression (GSR) was not included in the pre-registered analysis plan. However, since pre-registration, we have been persuaded by recent publications suggesting that GSR optimizes predicting individual differences in behavior from functional connectivity estimates (Finn & Bandettini, 2021; Li et al., 2019). We report the results without global signal regression in the Supplemental Materials. The conclusions regarding the DMN remain the same across methods, with some differences in the hippocampal and CBPM analyses noted in the Results and Supplemental Materials.

## References

Anderson, J. S., Ferguson, M. A., Lopez-Larson, M., & Yurgelun-Todd, D. (2011). Reproducibility of single-subject functional connectivity measurements. AJNR. American Journal of Neuroradiology, 32(3), 548–555.

Andrews-Hanna, J. R., Reidler, J. S., Sepulcre, J., Poulin, R., & Buckner, R. L. (2010). Functional-anatomic fractionation of the brain’s default network. Neuron, 65(4), 550–562.

Avery, E. W., Yoo, K., Rosenberg, M. D., Greene, A. S., Gao, S., Na, D. L., Scheinost, D., Constable, T. R., & Chun, M. M. (2020). Distributed Patterns of Functional Connectivity Predict Working Memory Performance in Novel Healthy and Memory-impaired Individuals. Journal of Cognitive Neuroscience, 32(2), 241–255.

Barnett, A. J., Reilly, W., Dimsdale-Zucker, H. R., Mizrak, E., Reagh, Z., & Ranganath, C. (2021). Intrinsic connectivity reveals functionally distinct cortico-hippocampal networks in the human brain. PLoS Biology, 19(6), e3001275.

Beaty, R. E., Kenett, Y. N., Christensen, A. P., Rosenberg, M. D., Benedek, M., Chen, Q., Fink, A., Qiu, J., Kwapil, T. R., Kane, M. J., & Silvia, P. J. (2018). Robust prediction of individual creative ability from brain functional connectivity. Proceedings of the National Academy of Sciences of the United States of America, 115(5), 1087–1092.

Bird, C. M., & Burgess, N. (2008). The hippocampus and memory: insights from spatial processing. Nature Reviews. Neuroscience, 9(3), 182–194.

Buckner, R. L., Andrews-Hanna, J. R., & Schacter, D. L. (2008). The brain’s default network: anatomy, function, and relevance to disease. Annals of the New York Academy of Sciences, 1124, 1–38.

Buckner, R. L., & DiNicola, L. M. (2019). The brain’s default network: updated anatomy, physiology and evolving insights. Nature Reviews Neuroscience. 10.1038/s41583-019-0212-7

Cabeza, R., & Moscovitch, M. (2013). Memory Systems, Processing Modes, and Components: Functional Neuroimaging Evidence. Perspectives on Psychological Science: A Journal of the Association for Psychological Science, 8(1), 49–55.

Clark, I. A., & Maguire, E. A. (2020). Do questionnaires reflect their purported cognitive functions? Cognition, 195, 104114.

Cooper, R. A., Kurkela, K. A., Davis, S. W., & Ritchey, M. (2021). Mapping the organization and dynamics of the posterior medial network during movie watching. NeuroImage, 236, 118075.

Cooper, R. A., & Ritchey, M. (2022). Patterns of episodic content and specificity predicting subjective memory vividness. Memory & Cognition. 10.3758/s13421-022-01291-5

Corkin, S. (2002). What’s new with the amnesic patient H.M.? Nature Reviews Neuroscience, 3(2), 153–160.

Davis, S. W., Stanley, M. L., Moscovitch, M., & Cabeza, R. (2017). Resting-state networks do not determine cognitive function networks: a commentary on Campbell and Schacter (2016). Language, Cognition and Neuroscience, 32(6), 669–673.

Diana, R. A., Yonelinas, A. P., & Ranganath, C. (2007). Imaging recollection and familiarity in the medial temporal lobe: a three-component model. Trends in Cognitive Sciences, 11(9), 379–386.

Elliott, M. L., Knodt, A. R., Cooke, M., Kim, M. J., Melzer, T. R., Keenan, R., Ireland, D., Ramrakha, S., Poulton, R., Caspi, A., Moffitt, T. E., & Hariri, A. R. (2019). General functional connectivity: Shared features of resting-state and task fMRI drive reliable and heritable individual differences in functional brain networks. NeuroImage, 189, 516–532.

Esteban, O., Birman, D., Schaer, M., Koyejo, O. O., Poldrack, R. A., & Gorgolewski, K. J. (2017). MRIQC: Advancing the automatic prediction of image quality in MRI from unseen sites. PloS One, 12(9), e0184661.

Esteban, O., Blair, R., Markiewicz, C. J., Berleant, S. L., Moodie, C., Ma, F., Isik, A. I., Erramuzpe, A., Kent, M., Goncalves, M., & Others. (2018). fmriprep. Software: Practice & Experience.

Esteban, O., Markiewicz, C. J., Blair, R. W., Moodie, C. A., Isik, A. I., Erramuzpe, A., Kent, J. D., Goncalves, M., DuPre, E., Snyder, M., Oya, H., Ghosh, S. S., Wright, J., Durnez, J., Poldrack, R. A., & Gorgolewski, K. J. (2019). fMRIPrep: a robust preprocessing pipeline for functional MRI. Nature Methods, 16(1), 111–116.

Finn, E. S., & Bandettini, P. A. (2021). Movie-watching outperforms rest for functional connectivity-based prediction of behavior. NeuroImage, 235, 117963.

Finn, E. S., Shen, X., Scheinost, D., Rosenberg, M. D., Huang, J., Chun, M. M., Papademetris, X., & Constable, R. T. (2015). Functional connectome fingerprinting: identifying individuals using patterns of brain connectivity. Nature Neuroscience, 18(11), 1664–1671.

Gohel, D., & Skintzos, P. (2023). flextable: Functions for Tabular Reporting. https://CRAN.R-project.org/package=flextable

Gordon, E. M., Laumann, T. O., Adeyemo, B., & Petersen, S. E. (2017). Individual Variability of the System-Level Organization of the Human Brain. Cerebral Cortex, 27(1), 386–399.

Gordon, E. M., Laumann, T. O., Gilmore, A. W., Newbold, D. J., Greene, D. J., Berg, J. J., Ortega, M., Hoyt-Drazen, C., Gratton, C., Sun, H., Hampton, J. M., Coalson, R. S., Nguyen, A. L., McDermott, K. B., Shimony, J. S., Snyder, A. Z., Schlaggar, B. L., Petersen, S. E., Nelson, S. M., & Dosenbach, N. U. F. (2017). Precision Functional Mapping of Individual Human Brains. Neuron, 95(4), 791–807.e7.

Gordon, E. M., & Nelson, S. M. (2021). Three types of individual variation in brain networks revealed by single-subject functional connectivity analyses. Current Opinion in Behavioral Sciences, 40, 79–86.

Gratton, C., Laumann, T. O., Nielsen, A. N., Greene, D. J., Gordon, E. M., Gilmore, A. W., Nelson, S. M., Coalson, R. S., Snyder, A. Z., Schlaggar, B. L., Dosenbach, N. U. F., & Petersen, S. E. (2018). Functional Brain Networks Are Dominated by Stable Group and Individual Factors, Not Cognitive or Daily Variation. Neuron, 98(2), 439–452.e5.

Greene, A. S., Gao, S., Noble, S., Scheinost, D., & Constable, R. T. (2020). How Tasks Change Whole-Brain Functional Organization to Reveal Brain-Phenotype Relationships. Cell Reports, 32(8), 108066.

Greene, A. S., Gao, S., Scheinost, D., & Constable, R. T. (2018). Task-induced brain state manipulation improves prediction of individual traits. Nature Communications, 9(1), 1–13.

Hsu, W.-T., Rosenberg, M. D., Scheinost, D., Constable, R. T., & Chun, M. M. (2018). Resting-state functional connectivity predicts neuroticism and extraversion in novel individuals. Social Cognitive and Affective Neuroscience, 13(2), 224–232.

Jääskeläinen, I. P., Sams, M., Glerean, E., & Ahveninen, J. (2021). Movies and narratives as naturalistic stimuli in neuroimaging. NeuroImage, 224, 117445.

Jangraw, D. C., Gonzalez-Castillo, J., Handwerker, D. A., Ghane, M., Rosenberg, M. D., Panwar, P., & Bandettini, P. A. (2018). A functional connectivity-based neuromarker of sustained attention generalizes to predict recall in a reading task. NeuroImage, 166, 99–109.

King, D. R., de Chastelaine, M., Elward, R. L., Wang, T. H., & Rugg, M. D. (2015). Recollection-related increases in functional connectivity predict individual differences in memory accuracy. Journal of Neuroscience, 35(4), 1763–1772.

Kong, R., Li, J., Orban, C., Sabuncu, M. R., Liu, H., Schaefer, A., Sun, N., Zuo, X.-N., Holmes, A. J., Eickhoff, S. B., & Yeo, B. T. T. (2019). Spatial Topography of Individual-Specific Cortical Networks Predicts Human Cognition, Personality, and Emotion. Cerebral Cortex, 29(6), 2533–2551.

Kurkela, K. A., Cooper, R. A., Ryu, E., & Ritchey, M. (2022). Integrating Region- and Network-level Contributions to Episodic Recollection Using Multilevel Structural Equation Modeling. Journal of Cognitive Neuroscience, 34(12), 2341–2359.

Laumann, T. O., Gordon, E. M., Adeyemo, B., Snyder, A. Z., Joo, S. J., Chen, M.-Y., Gilmore, A. W., McDermott, K. B., Nelson, S. M., Dosenbach, N. U. F., Schlaggar, B. L., Mumford, J. A., Poldrack, R. A., & Petersen, S. E. (2015). Functional System and Areal Organization of a Highly Sampled Individual Human Brain. Neuron, 87(3), 657–670.

Lee, H., Bellana, B., & Chen, J. (2020). What can narratives tell us about the neural bases of human memory? Current Opinion in Behavioral Sciences, 32, 111–119.

LePort, A. K. R., Mattfeld, A. T., Dickinson-Anson, H., Fallon, J. H., Stark, C. E. L., Kruggel, F., Cahill, L., & McGaugh, J. L. (2012). Behavioral and neuroanatomical investigation of Highly Superior Autobiographical Memory (HSAM). Neurobiology of Learning and Memory, 98(1), 78–92.

Li, J., Kong, R., Liégeois, R., Orban, C., Tan, Y., Sun, N., Holmes, A. J., Sabuncu, M. R., Ge, T., & Yeo, B. T. T. (2019). Global signal regression strengthens association between resting-state functional connectivity and behavior. NeuroImage, 196, 126–141.

Lin, Q., Yoo, K., Shen, X., Constable, T. R., & Chun, M. M. (2021). Functional Connectivity during Encoding Predicts Individual Differences in Long-Term Memory. Journal of Cognitive Neuroscience, 1–18.

Marek, S., Tervo-Clemmens, B., Calabro, F. J., Montez, D. F., Kay, B. P., Hatoum, A. S., Donohue, M. R., Foran, W., Miller, R. L., Hendrickson, T. J., Malone, S. M., Kandala, S., Feczko, E., Miranda-Dominguez, O., Graham, A. M., Earl, E. A., Perrone, A. J., Cordova, M., Doyle, O., … Dosenbach, N. U. F. (2022). Reproducible brain-wide association studies require thousands of individuals. Nature, 603(7902), 654–660.

Matijevic, S., Andrews-Hanna, J. R., Wank, A. A., Ryan, L., & Grilli, M. D. (2022). Individual differences in the relationship between episodic detail generation and resting state functional connectivity vary with age. Neuropsychologia, 166, 108138.

Moscovitch, M., Cabeza, R., Winocur, G., & Nadel, L. (2016). Episodic Memory and Beyond: The Hippocampus and Neocortex in Transformation. Annual Review of Psychology, 67, 105–134.

Murphy, K., Birn, R. M., Handwerker, D. A., Jones, T. B., & Bandettini, P. A. (2009). The impact of global signal regression on resting state correlations: are anti-correlated networks introduced? NeuroImage, 44(3), 893–905.

Murphy, K., & Fox, M. D. (2017). Towards a consensus regarding global signal regression for resting state functional connectivity MRI. NeuroImage, 154, 169–173.

Ngo, C. T., Michelmann, S., Olson, I. R., & Newcombe, N. S. (2021). Pattern separation and pattern completion: Behaviorally separable processes? Memory & Cognition, 49(1), 193–205.

Palombo, D. J., Williams, L. J., Abdi, H., & Levine, B. (2013). The survey of autobiographical memory (SAM): a novel measure of trait mnemonics in everyday life. Cortex; a Journal Devoted to the Study of the Nervous System and Behavior, 49(6), 1526–1540.

Parker, E. S., Cahill, L., & McGaugh, J. L. (2006). A case of unusual autobiographical remembering. Neurocase, 12(1), 35–49.

Petrican, R., Palombo, D. J., Sheldon, S., & Levine, B. (2020). The Neural Dynamics of Individual Differences in Episodic Autobiographical Memory. eNeuro, 7(2). 10.1523/ENEURO.0531-19.2020

Power, J. D., Barnes, K. A., Snyder, A. Z., Schlaggar, B. L., & Petersen, S. E. (2012). Spurious but systematic correlations in functional connectivity MRI networks arise from subject motion. NeuroImage, 59(3), 2142–2154.

Power, J. D., Mitra, A., Laumann, T. O., Snyder, A. Z., Schlaggar, B. L., & Petersen, S. E. (2014). Methods to detect, characterize, and remove motion artifact in resting state fMRI. NeuroImage, 84, 320–341.

Przeździk, I., Faber, M., Fernández, G., Beckmann, C. F., & Haak, K. V. (2019). The functional organisation of the hippocampus along its long axis is gradual and predicts recollection. Cortex; a Journal Devoted to the Study of the Nervous System and Behavior, 119, 324–335.

R Core Team. (2022). R: A Language and Environment for Statistical Computing. R Foundation for Statistical Computing. https://www.R-project.org/

Reagh, Z. M., Delarazan, A. I., Garber, A., & Ranganath, C. (2020). Aging alters neural activity at event boundaries in the hippocampus and Posterior Medial network. Nature Communications, 11(1), 3980.

Richter, A., Soch, J., Kizilirmak, J. M., Fischer, L., Schütze, H., Assmann, A., Behnisch, G., Feldhoff, H., Knopf, L., Raschick, M., Schult, A., Seidenbecher, C. I., Yakupov, R., Düzel, E., & Schott, B. H. (2023). Single-value scores of memory-related brain activity reflect dissociable neuropsychological and anatomical signatures of neurocognitive aging. Human Brain Mapping, 44(8), 3283–3301.

Ritchey, M., & Cooper, R. A. (2020). Deconstructing the Posterior Medial Episodic Network. Trends in Cognitive Sciences. 10.1016/j.tics.2020.03.006

Ritchey, M., Montchal, M. E., Yonelinas, A. P., & Ranganath, C. (2015). Delay-dependent contributions of medial temporal lobe regions to episodic memory retrieval. eLife, 4. 10.7554/eLife.05025

Rosenberg, M. D., Finn, E. S., Scheinost, D., Papademetris, X., Shen, X., Constable, R. T., & Chun, M. M. (2016). A neuromarker of sustained attention from whole-brain functional connectivity. Nature Neuroscience, 19(1), 165–171.

Rugg, M. D., & Vilberg, K. L. (2013). Brain networks underlying episodic memory retrieval. Current Opinion in Neurobiology, 23(2), 255–260.

Satterthwaite, T. D., Ciric, R., Roalf, D. R., Davatzikos, C., Bassett, D. S., & Wolf, D. H. (2019). Motion artifact in studies of functional connectivity: Characteristics and mitigation strategies. Human Brain Mapping, 40(7), 2033–2051.

Satterthwaite, T. D., Elliott, M. A., Gerraty, R. T., Ruparel, K., Loughead, J., Calkins, M. E., Eickhoff, S. B., Hakonarson, H., Gur, R. C., Gur, R. E., & Wolf, D. H. (2013). An improved framework for confound regression and filtering for control of motion artifact in the preprocessing of resting-state functional connectivity data. NeuroImage, 64, 240–256.

Schaefer, A., Kong, R., Gordon, E. M., Laumann, T. O., Zuo, X.-N., Holmes, A. J., Eickhoff, S. B., & Yeo, B. T. T. (2018). Local-Global Parcellation of the Human Cerebral Cortex from Intrinsic Functional Connectivity MRI. Cerebral Cortex, 28(9), 3095–3114.

Seitzman, B. A., Gratton, C., Laumann, T. O., Gordon, E. M., Adeyemo, B., Dworetsky, A., Kraus, B. T., Gilmore, A. W., Berg, J. J., Ortega, M., Nguyen, A., Greene, D. J., McDermott, K. B., Nelson, S. M., Lessov-Schlaggar, C. N., Schlaggar, B. L., Dosenbach, N. U. F., & Petersen, S. E. (2019). Trait-like variants in human functional brain networks. Proceedings of the National Academy of Sciences of the United States of America. 10.1073/pnas.1902932116

Setton, R., Lockrow, A. W., Turner, G. R., & Spreng, R. N. (2022). Troubled past: A critical psychometric assessment of the self-report Survey of Autobiographical Memory (SAM). Behavior Research Methods, 54(1), 261–286.

Setton, R., Mwilambwe-Tshilobo, L., Sheldon, S., Turner, G. R., & Spreng, R. N. (2022). Hippocampus and temporal pole functional connectivity is associated with age and individual differences in autobiographical memory. Proceedings of the National Academy of Sciences of the United States of America, 119(41), e2203039119.

Shafto, M. A., Tyler, L. K., Dixon, M., Taylor, J. R., Rowe, J. B., Cusack, R., Calder, A. J., Marslen-Wilson, W. D., Duncan, J., Dalgleish, T., Henson, R. N., Brayne, C., Matthews, F. E., & Cam-CAN. (2014). The Cambridge Centre for Ageing and Neuroscience (Cam-CAN) study protocol: a cross-sectional, lifespan, multidisciplinary examination of healthy cognitive ageing. BMC Neurology, 14, 204.

Sheldon, S., Farb, N., Palombo, D. J., & Levine, B. (2016). Intrinsic medial temporal lobe connectivity relates to individual differences in episodic autobiographical remembering. Cortex; a Journal Devoted to the Study of the Nervous System and Behavior, 74, 206–216.

Shen, X., Finn, E. S., Scheinost, D., Rosenberg, M. D., Chun, M. M., Papademetris, X., & Constable, R. T. (2017). Using connectome-based predictive modeling to predict individual behavior from brain connectivity. Nature Protocols, 12(3), 506–518.

Sjoberg, D., Whiting, K., Curry, M., Lavery, J., & Larmarange, J. (2021). Reproducible Summary Tables with the gtsummary Package. The R Journal, 13(1), 570.

Sneve, M. H., Grydeland, H., Amlien, I. K., Langnes, E., Walhovd, K. B., & Fjell, A. M. (2017). Decoupling of large-scale brain networks supports the consolidation of durable episodic memories. NeuroImage, 153, 336–345.

Squire, L. R. (1992). Memory and the hippocampus: a synthesis from findings with rats, monkeys, and humans. Psychological Review, 99(2), 195–231.

Tambini, A., Ketz, N., & Davachi, L. (2010). Enhanced brain correlations during rest are related to memory for recent experiences. Neuron, 65(2), 280–290.

Taylor, J. R., Williams, N., Cusack, R., Auer, T., Shafto, M. A., Dixon, M., Tyler, L. K., Cam-Can, & Henson, R. N. (2017). The Cambridge Centre for Ageing and Neuroscience (Cam-CAN) data repository: Structural and functional MRI, MEG, and cognitive data from a cross-sectional adult lifespan sample. NeuroImage, 144(Pt B), 262–269.

Touroutoglou, A., Andreano, J. M., Barrett, L. F., & Dickerson, B. C. (2015). Brain network connectivity-behavioral relationships exhibit trait-like properties: Evidence from hippocampal connectivity and memory. Hippocampus, 25(12), 1591–1598.

Unsworth, N. (2019). Individual differences in long-term memory. Psychological Bulletin, 145(1), 79–139.

van Buuren, M., Wagner, I. C., & Fernández, G. (2019). Functional network interactions at rest underlie individual differences in memory ability. Learning & Memory, 26(1), 9–19.

Wang, L., LaViolette, P., O’Keefe, K., Putcha, D., & Bakkour, A. (2010). Intrinsic connectivity between the hippocampus and posteromedial cortex predicts memory performance in cognitively intact older individuals. NeuroImage. https://www.sciencedirect.com/science/article/pii/S1053811910002144

Wang, L., Negreira, A., LaViolette, P., Bakkour, A., Sperling, R. A., & Dickerson, B. C. (2010). Intrinsic interhemispheric hippocampal functional connectivity predicts individual differences in memory performance ability. Hippocampus, 20(3), 345–351.

Wang, Z., Goerlich, K. S., Ai, H., Aleman, A., Luo, Y.-J., & Xu, P. (2021). Connectome-Based Predictive Modeling of Individual Anxiety. Cerebral Cortex, 31(6), 3006–3020.

Weschler, C. J. (1999). Weschler Memory Scale (Third UK edition).

Whitfield-Gabrieli, S., & Nieto-Castanon, A. (2012). Conn: a functional connectivity toolbox for correlated and anticorrelated brain networks. Brain Connectivity, 2(3), 125–141.

Wickham, H. (2016). ggplot2: Elegant Graphics for Data Analysis. Springer-Verlag New York. https://ggplot2.tidyverse.org

Wing, E. A., Marsh, E. J., & Cabeza, R. (2013). Neural correlates of retrieval-based memory enhancement: an fMRI study of the testing effect. Neuropsychologia, 51(12), 2360–2370.

Yeo, B. T. T., Krienen, F. M., Sepulcre, J., Sabuncu, M. R., Lashkari, D., Hollinshead, M., Roffman, J. L., Smoller, J. W., Zöllei, L., Polimeni, J. R., Fischl, B., Liu, H., & Buckner, R. L. (2011). The organization of the human cerebral cortex estimated by intrinsic functional connectivity. Journal of Neurophysiology, 106(3), 1125–1165.

Zhao, W., Makowski, C., Hagler, D. J., Garavan, H. P., Thompson, W. K., Greene, D. J., Jernigan, T. L., & Dale, A. M. (2023). Task fMRI paradigms may capture more behaviorally relevant information than resting-state functional connectivity. NeuroImage, 270, 119946.

